## Supplementary Fig. 1 for "Purine and pyrimidine synthesis differently affect the strength of the inoculum effect for aminoglycoside and β-lactam antibiotics"

### Supplementary Figures and Figure Legends

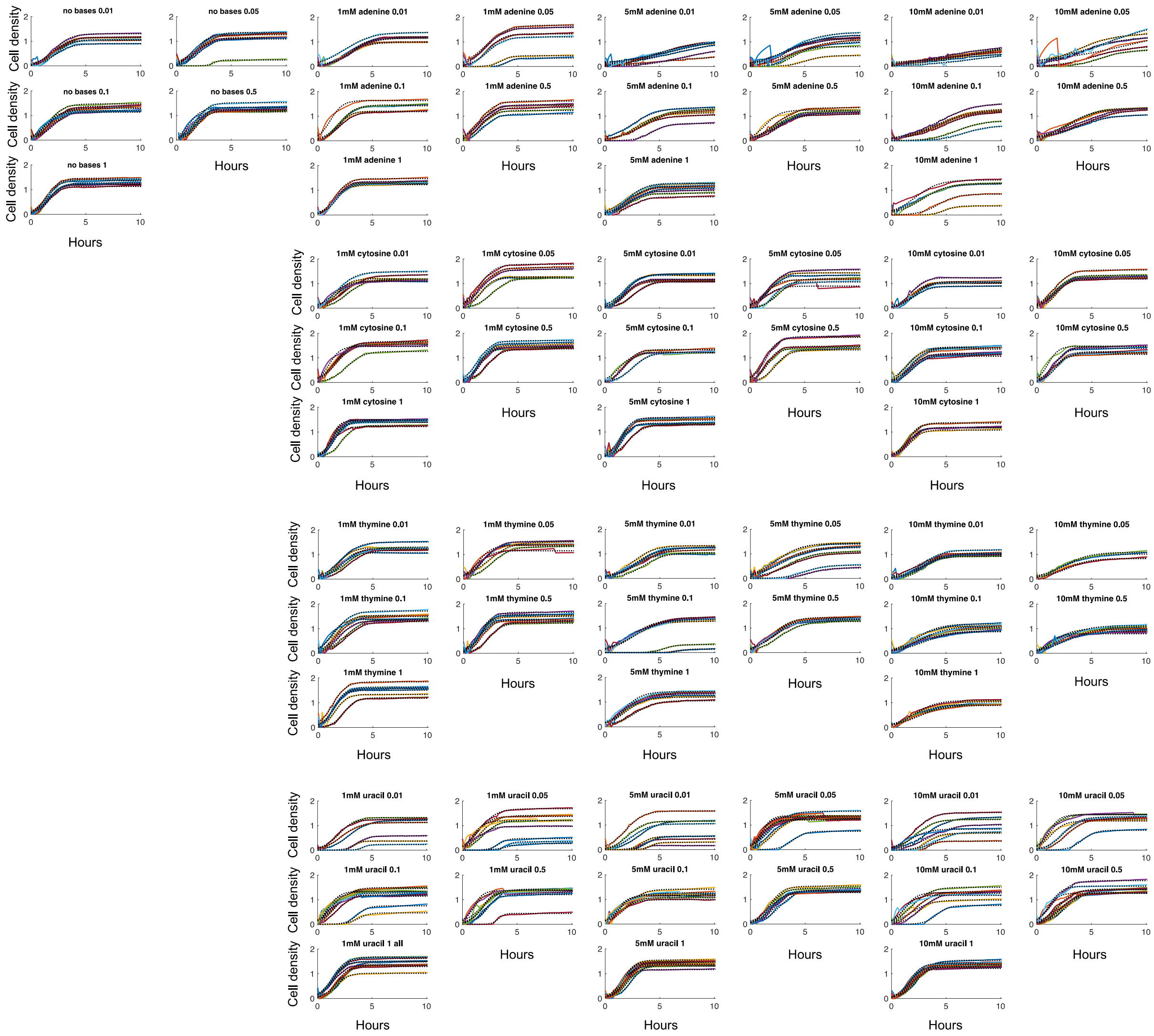


**Supplementary Figure 1: Growth curves of *E. coli* grown in M9 medium with different concentrations of nitrogenous bases as indicated.** The numbers at the top of each plot indicate the percentage of casamino acids (e.g., 0.01 = 0.01%). Colored lines = experimental data. Black dotted lines = growth curve fit using a logistic equation.
