## Supplementary Fig. 2 for "Purine and pyrimidine synthesis differently affect the strength of the inoculum effect for aminoglycoside and β-lactam antibiotics"

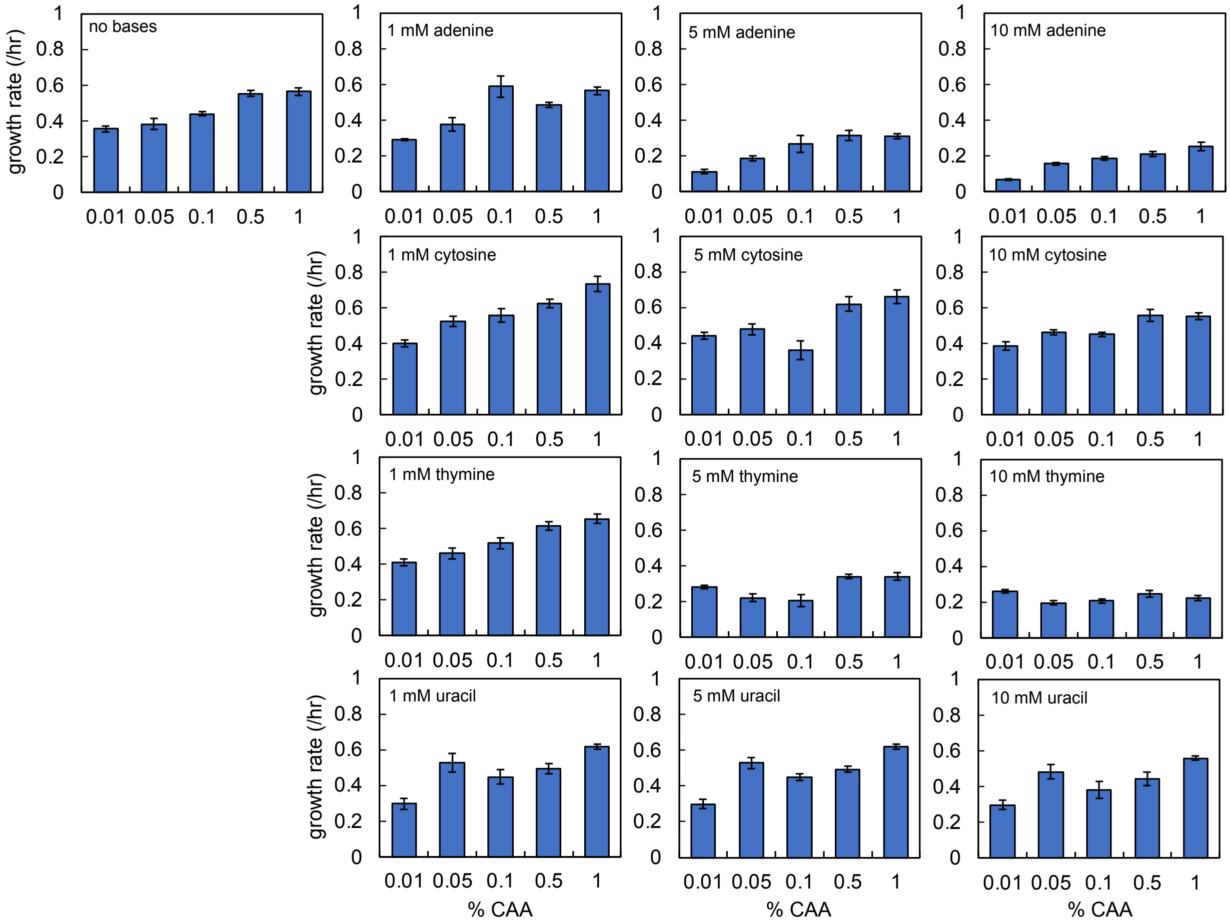


**Supplementary Figure 2: Raw growth rates for *E. coli* grown in M9 medium with different concentrations of nitrogenous bases as indicated on the plot**. % CAA = percentage of casamino acids. Error bars = SEM. Bars = average from ≥ 4 biological replicates
