## Supplementary Fig. 3 for "Purine and pyrimidine synthesis differently affect the strength of the inoculum effect for aminoglycoside and β-lactam antibiotics"

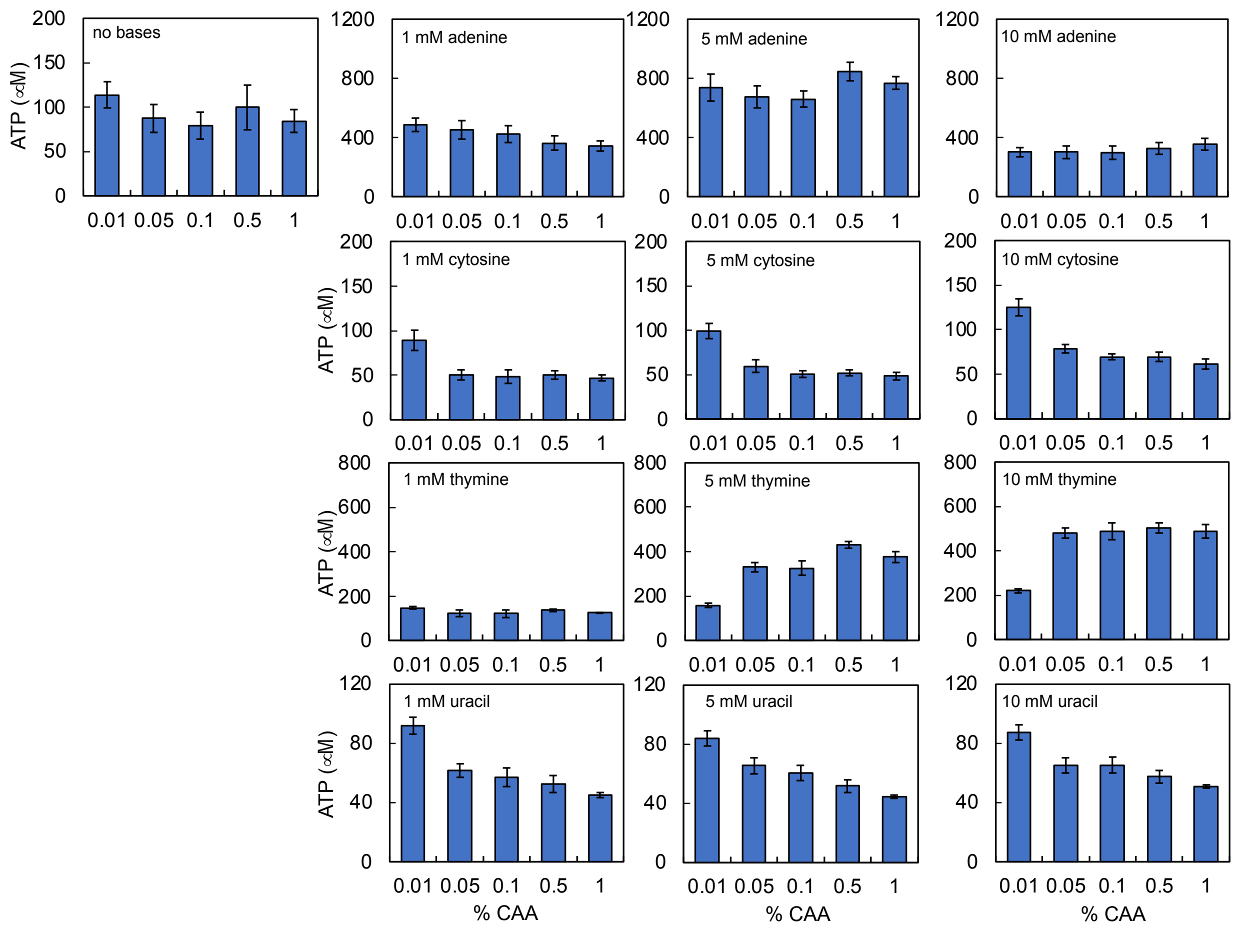


**Supplementary Figure 3: Raw [ATP] for *E. coli* grown in M9 medium with different concentrations of nitrogenous bases as indicated.** % CAA = percentage of casamino acids. Error bars = SEM. Bars = average from 4 biological replicates
