## Supplementary Fig. 4 for "Purine and pyrimidine synthesis differently affect the strength of the inoculum effect for aminoglycoside and β-lactam antibiotics"

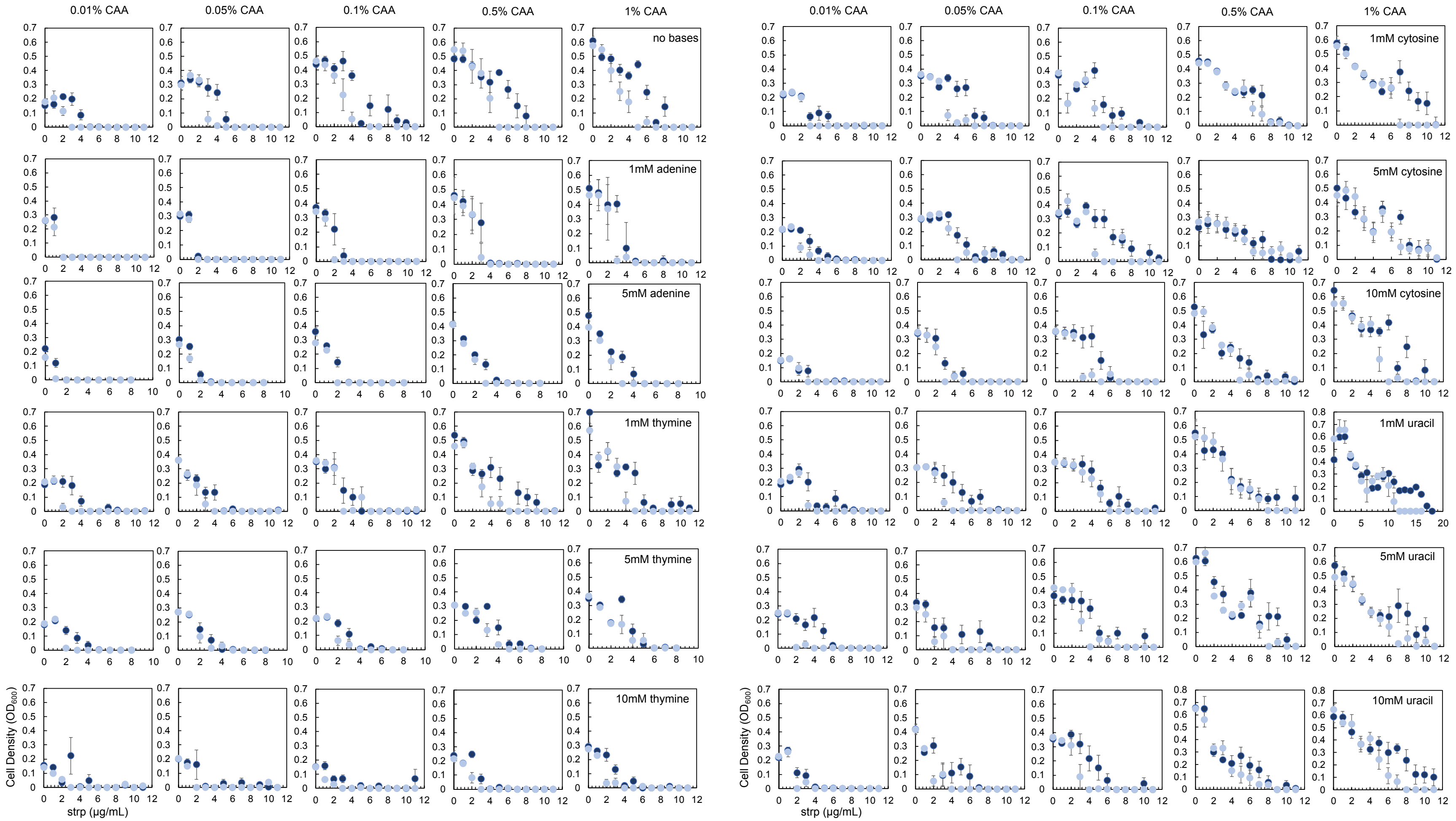


**Supplementary Figure 4: Raw MIC data for *E. coli* grown in streptomycin (strp).** % CAA = percentage of casamino acids. Error bars = SEM. Data corresponds to Fig. 3A. Each data point is the average of ≥ 5 biological replicates. Dark blue = high density; light blue = low density.
