## Supplementary Fig. 5 for "Purine and pyrimidine synthesis differently affect the strength of the inoculum effect for aminoglycoside and β-lactam antibiotics"

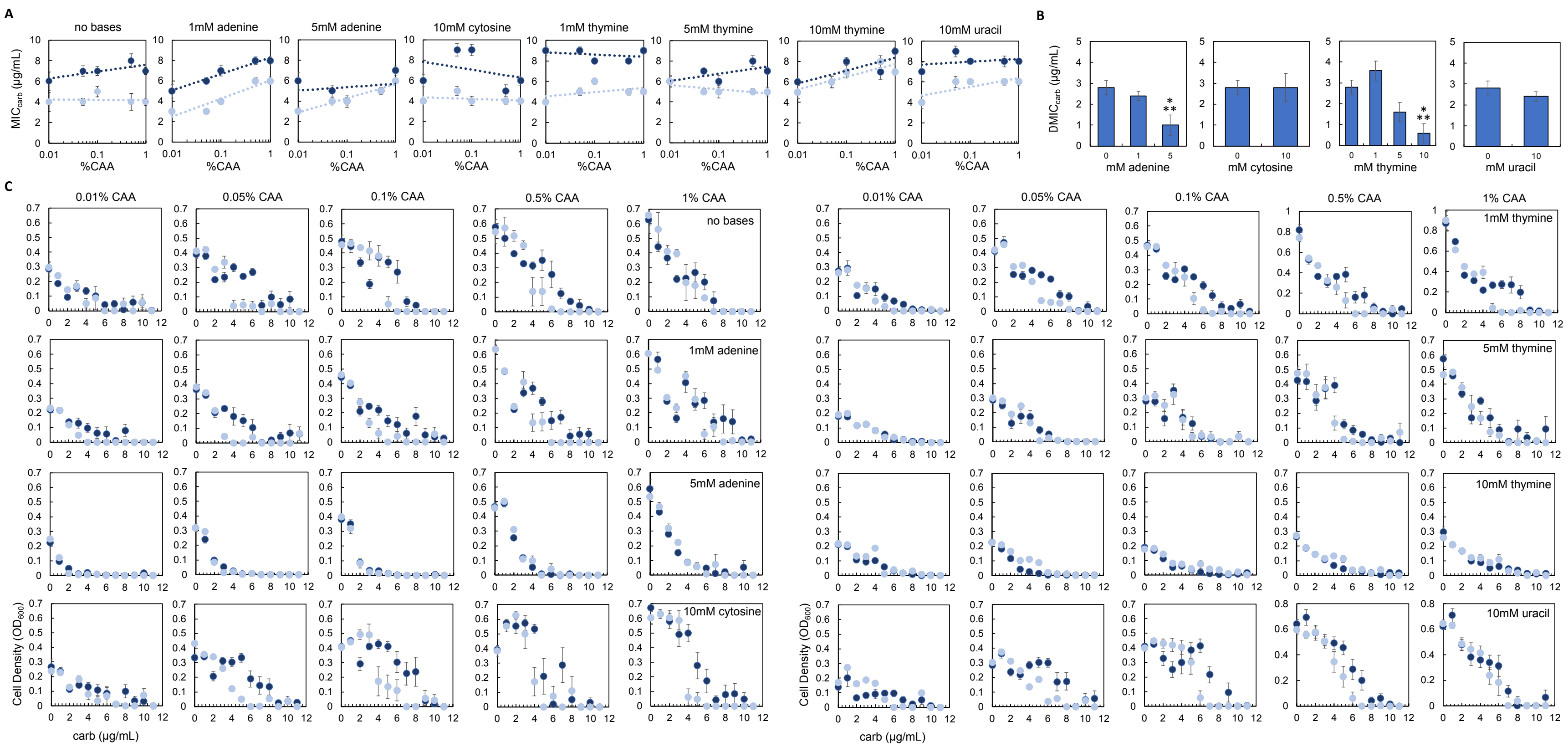


**Supplementary Figure 5: MIC data for *E. coli* grown in carbenicillin.**

A) MIC of carbenicillin (carb) as a function of the percentage of casamino acids. % CAA = percentage of casamino acids. Dark blue = high initial density, light blue = low initial density. For all panels error bars = SEM and each data point is the average of ≥ 5 biological replicates.

B) Average $\Delta MIC_{carb}$ for each growth condition. Average plotted from 5 different % CAA each consisting of ≥ 5 biological replicates. Error bars = SEM. * different than no nitrogenous base control ($P \leq 0.03$, two-tailed t-test); ** not different than zero ($P \geq 0.071$, one-tailed t-test).

C) Raw data for panel A. Dark blue = high initial density, light blue = low initial density. Each data point is averaged from ≥ 5 biological replicates.
