## Supplementary Fig. 7 for "Purine and pyrimidine synthesis differently affect the strength of the inoculum effect for aminoglycoside and β-lactam antibiotics"

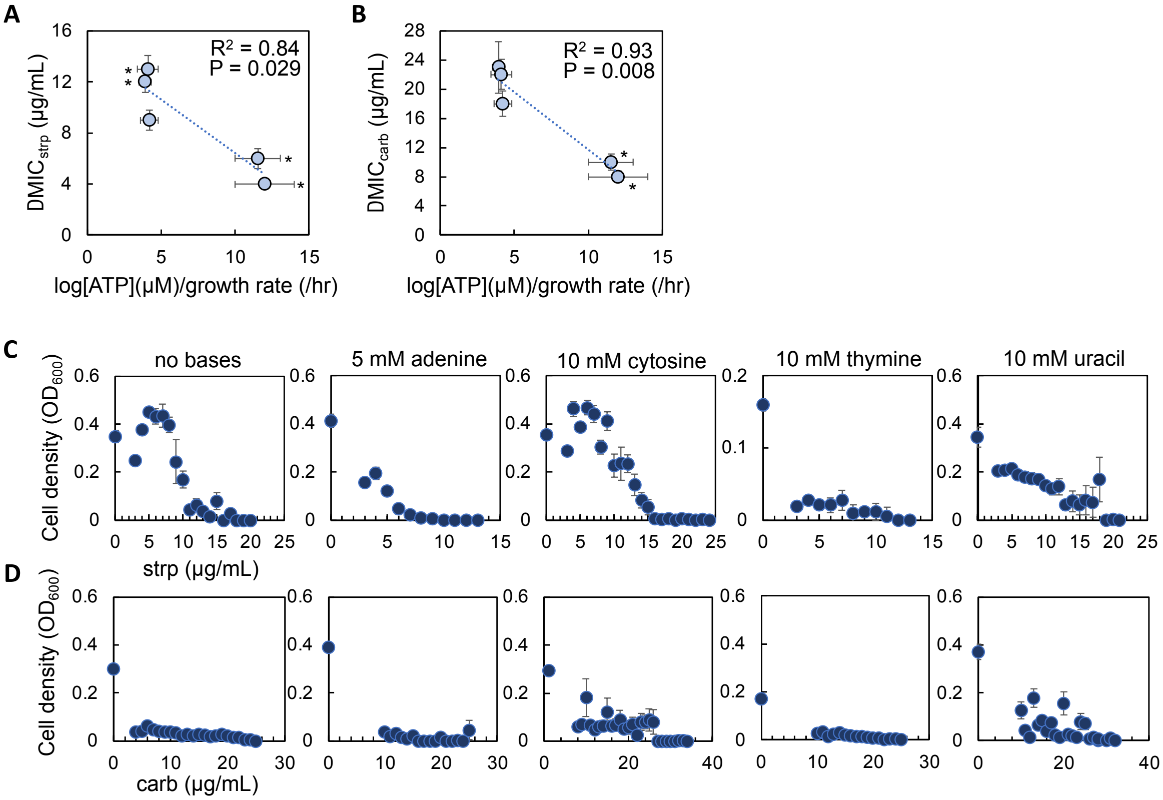


**Supplementary Figure 7: The relationship between log[ATP]/growth rate and ΔMIC when a higher density of bacteria is used ($2.00 \times 10^7$ CFU/mL +/- $4.50 \times 10^6$).**

- **A)** ΔMIC of streptomycin (strp) as a function of log[ATP]/growth rate. $R^2$ and P value from a linear regression.* significantly different than no nitrogenous base control ($P \leq 0.002$, two-tailed t-test). log[ATP]/growth from Fig. 2C. Average plotted from ≥ 6 biological replicates. Error bars = SEM. WLS: $R^2 = 0.89$, $P = 0.015$, Deming regression: $P = 0.0290$. For all panels, ΔMIC was only measured using medium with 0.1% CAA.
- **B)** ΔMIC of carbenicillin (carb) as a function of log[ATP]/growth rate. $R^2$ and P value from a linear regression.* significantly different than no nitrogenous base control ($P \leq 0.001$, two-tailed t-test); log[ATP]/growth from Fig. 2C. Average plotted from ≥ 4 biological replicates. Error bars = SEM. WLS: $R^2 = 0.83$, $P = 0.032$, Deming regression: $P = 0.033$.
- **C)** Raw MIC data for experiments with a higher density of bacteria challenged with strp. Average from ≥ 4 biological replicates. Error bars = SEM.
- **D)** Raw MIC data for experiments with a higher density of bacteria challenged with carb. Average from ≥ 4 biological replicates. Error bars = SEM.
