## Supplementary Fig. 8 for "Purine and pyrimidine synthesis differently affect the strength of the inoculum effect for aminoglycoside and β-lactam antibiotics"

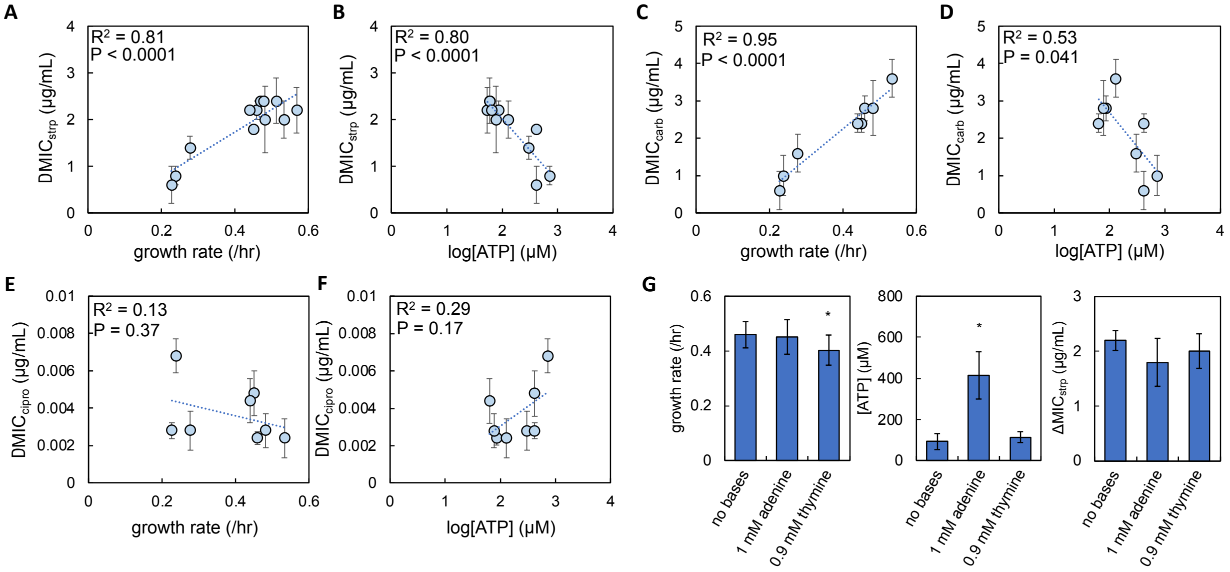


**Supplementary Figure 8: Regression analysis between ΔMIC, and growth rate, log[ATP], and carrying capacity for each antibiotic.**

- **A)** The relationship between ΔMIC of streptomycin (strp) and growth rate (/hr). For panels A-F, data from Fig. 2 (growth and [ATP]) and 3 (ΔMIC). $R^2$ and P value from a linear regression. Error bars = SEM.
- **B)** The relationship between ΔMIC of strp and log[ATP].
- **C)** The relationship between ΔMIC of carbenicillin (carb) and growth rate.
- **D)** The relationship between ΔMIC of carb and log[ATP].
- **E)** The relationship between ΔMIC of ciprofloxacin (cipro) and growth rate.
- **F)** The relationship between ΔMIC of cipro and log[ATP].
- **G)** The effect of changing only growth rate or [ATP] using nitrogenous bases on the ΔMIC of strp. Errors = SEM. Average [ATP] from four biological replicates. Average growth rate from ≥ 6 biological replicates. ΔMIC from ≥ 4 biological replicates. * = statistically different from no nitrogenous base control. Statistics using two-tailed t-test: growth rate (1 mM adenine vs control, P = 0.567; 0.9 mM thymine vs control, P = 0.031), [ATP] (1 mM adenine vs control, P < 0.001; 0.9 mM thymine vs control, P = 0.086), ΔMIC (1 mM adenine vs control, P = 0.48; 0.9 mM thymine vs control, P = 0.61).
