## Supplementary Fig. 9 for "Purine and pyrimidine synthesis differently affect the strength of the inoculum effect for aminoglycoside and β-lactam antibiotics"

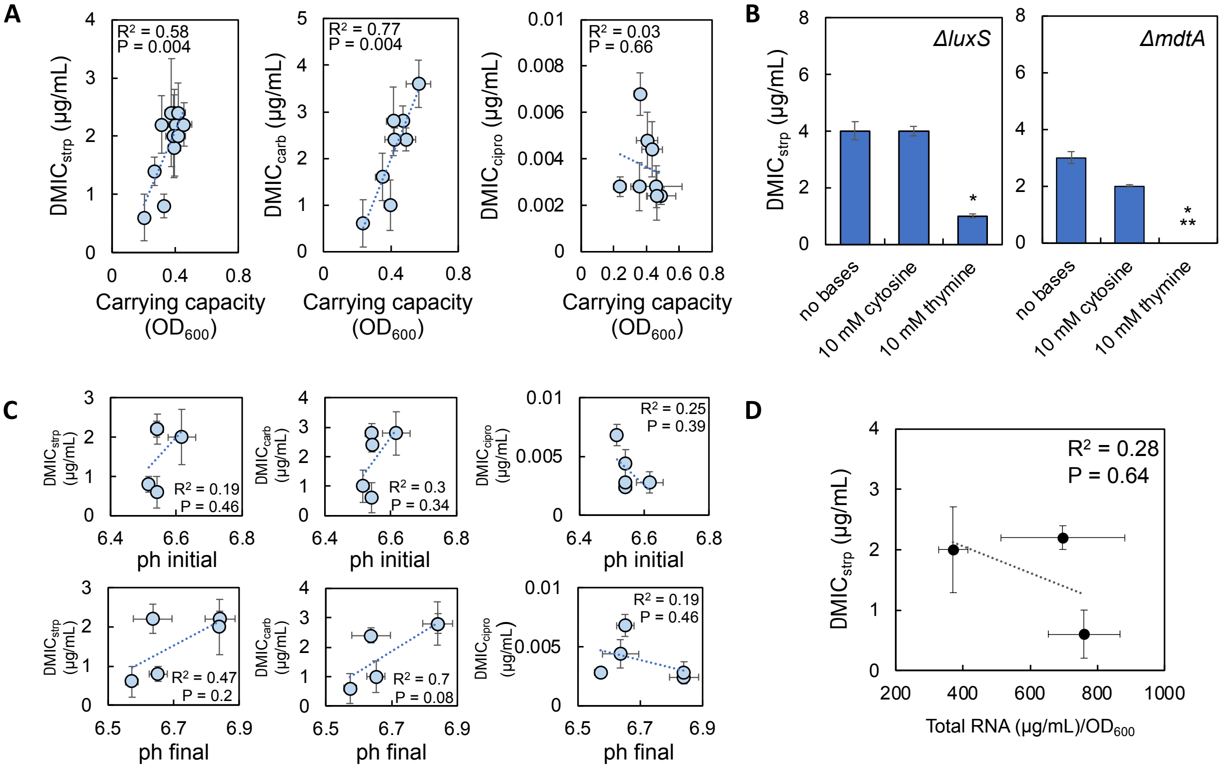

**Supplementary Figure 9: Alternative explanations for IE.**

**A)** The relationship between ΔMIC of strp (left), carb (center), and cipro (right) and carrying capacity of the growth medium (measured using $OD_{600}$ of bacteria grown in the absence of antibiotics; averaged from both high and low density populations).

**B)** ΔMIC of streptomycin (strp) for *E. coli* strains that lack quorum sensing (*luxS*) or efflux pumps (*mdtA*). All strains retained IE and supplementation with 10 mM thymine, but not 10 mM cytosine, reduced ΔMIC. Error bars = SEM. Average from ≥ 5 biological replicates. * indicates less than no nitrogenous base (no bases.) control ($P \leq 0.019$, two-tailed t-test). ** indicates not different than zero ($P = 0.104$, one-tailed t-test).

**C)** Linear regression between the pH of the growth medium before (top) and after (bottom) 24 hours of bacterial growth. The medium does not contain antibiotics. Error bars = SEM. pH averaged from 3 biological replicates. ΔMIC from Fig. 3.

**D)** The effect of adding 10 mM cytosine or 10 mM thymine on the concentration of rRNA in the cell. Error bars = SEM. Average from ≥ 2 biological replicates consisting of 3 technical replicates. ΔMIC of strp from Fig. 3. $P = 0.15$, ANOVA ($P > 0.16$, Tukey's HSD all comparisons).
