## Supplementary Fig. 10 for "Purine and pyrimidine synthesis differently affect the strength of the inoculum effect for aminoglycoside and β-lactam antibiotics"

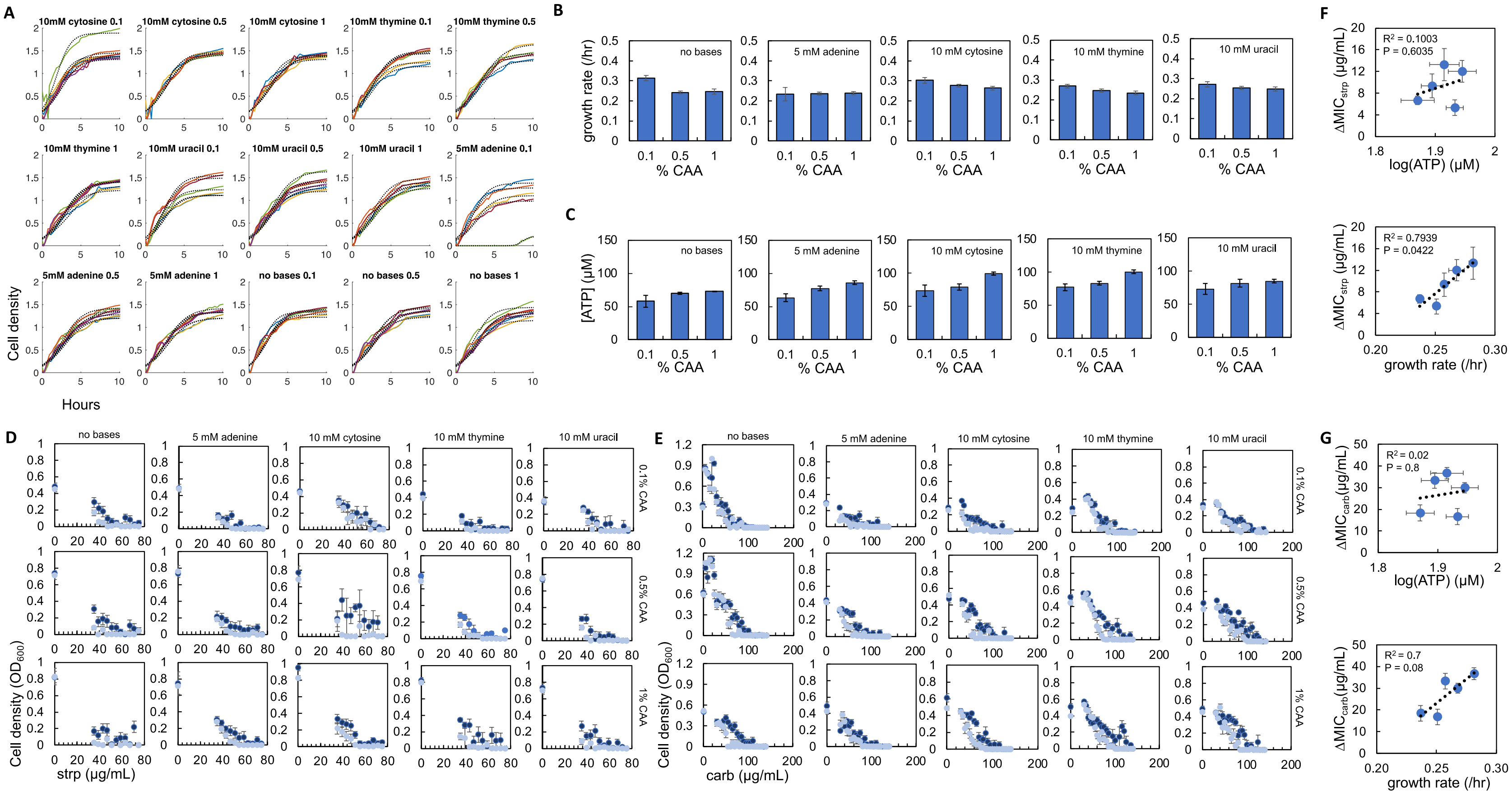


**Supplementary Figure 10: Raw data for *P. aeruginosa*.**

**A)** Growth curves of *P. aeruginosa* grown in M9 medium with different concentrations of nitrogenous bases as indicated**.** The numbers at the top of each plot indicate the percentage of casamino acids (0.1 = 0.1% CAA). Colored lines = experimental data. Black dotted lines = growth curve fit using logistic equation.

**B)** Average growth rate for each percentage of casamino acids (CAA) used. Error bars = SEM. Average plotted from ≥ 3 biological replicates.

**C)** Average [ATP] for each % CAA and with, or without, nitrogenous bases as indicated. Error bars = SEM. Average plotted from ≥ 4 biological replicates.

**D)** Raw MIC data for *P. aeruginosa* grown in streptomycin (strp). Error bars = SEM. Average from ≥ 5 biological replicates. Dark blue = high density; light blue = low density.

**E)** Raw MIC data for *P. aeruginosa* grown in carbenicillin (carb). Error bars = SEM. Average from ≥ 5 biological replicates. Dark blue = high density; light blue = low density.

**F)** The relationship between ΔMIC of strp, [ATP] (top panel), and growth rate (bottom panel). ΔMIC, growth rate, and [ATP] from Fig. 4. $R^2$ and P value from a linear regression. Error bars = SEM.

**G)** The relationship between ΔMIC of carb, [ATP] (top panel), and growth rate (bottom panel). ΔMIC, growth rate, and [ATP] from Fig. 4. $R^2$ and P value from a linear regression. Error bars = SEM.
