## Supplementary Fig. 11 for "Purine and pyrimidine synthesis differently affect the strength of the inoculum effect for aminoglycoside and β-lactam antibiotics"

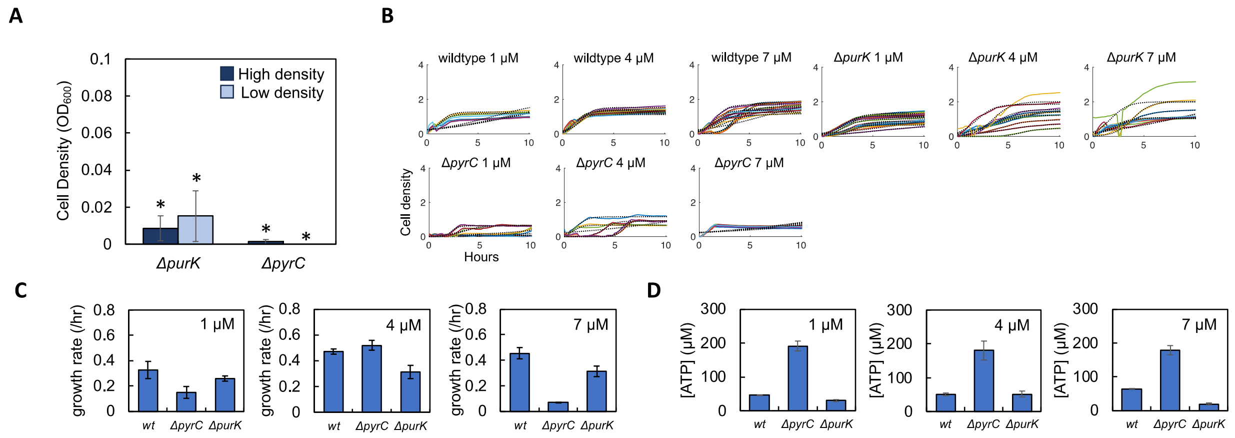


**Supplementary Figure 11: Raw data for growth, growth rates, and [ATP] for knockout strains.**

- **A)** Cell density ($OD_{600}$) of Δ*pyrC* and Δ*purK* knockout strains grown in M9 medium without exogenously supplemented nitrogenous bases. Average from three biological replicates of bacteria in medium with 0.01%, 0.05%, 0.1%, 0.5% and 1% casamino acids. Error bars = SEM. P > 0.16 when compared against zero (* = not different than zero, one-tailed t-test).
- **B)** Growth curves of *E. coli* knockout strains grown in M9 medium with different concentrations of equimolar nitrogenous bases as indicated in the plot. Colored lines = experimental data. Black dotted lines = growth curve fit.
- **C)** Raw growth rates. Concentration of equimolar nitrogenous bases is indicated on the plot. Error bars = SEM. Bars = average from ≥ 5 biological replicates.
- **D)** Raw [ATP] data. Concentration of equimolar nitrogenous bases is indicated on the plot. Error bars = SEM. Average from 3 biological replicates.
