## Supplementary Fig. 12 for "Purine and pyrimidine synthesis differently affect the strength of the inoculum effect for aminoglycoside and β-lactam antibiotics"

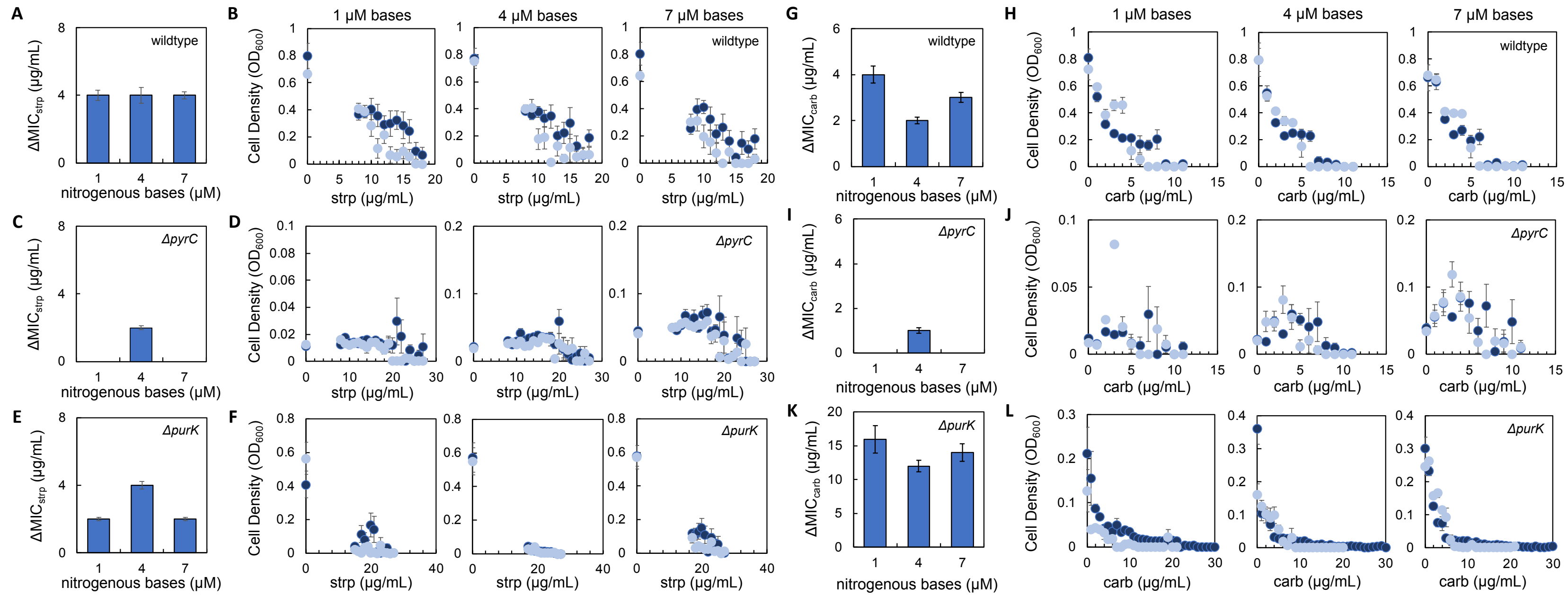


**Supplementary Figure 12: Raw MIC data for purine and pyrimidine synthesis knockout strains.**

- **A)** ΔMIC of streptomycin (strp) for wildtype *E. coli* as a function of the concentration of equimolar nitrogenous bases provided in the growth medium. Average from ≥ 5 biological replicates. For all panels, error bars = SEM.
- **B)** Raw data for panel A. For panels B, D, F, H, J, and L, dark blue = high initial density, light blue = low initial density.
- **C)** ΔMIC of strp for Δ*pyrC* as a function of the concentration of equimolar nitrogenous bases provided in the growth medium. Average from ≥ 5 biological replicates.
- **D)** Raw data for panel C.
- **E)** ΔMIC of strp for Δ*purK* as a function of the concentration of equimolar nitrogenous bases provided in the growth medium. Average from ≥ 5 biological replicates.
- **F)** Raw data for panel E.
- **G)** ΔMIC of carbenicillin (carb) for wildtype *E. coli* as a function of the concentration of equimolar nitrogenous bases provided in the growth medium. Average from ≥ 5 biological replicates.

**H)** Raw data for panel G.

**I)** ΔMIC of carb for Δ*pyrC* as a function of the concentration of equimolar nitrogenous bases provided in the growth medium. Average from ≥ 5 biological replicates.

**J)** Raw data for panel I.

**K)** ΔMIC of carb for Δ*purK* as a function of c equimolar nitrogenous bases provided in the growth medium. Average from ≥ 5 biological replicates.

**L)** Raw data for panel K.
