## Supplementary Fig. 13 for "Purine and pyrimidine synthesis differently affect the strength of the inoculum effect for aminoglycoside and β-lactam antibiotics"

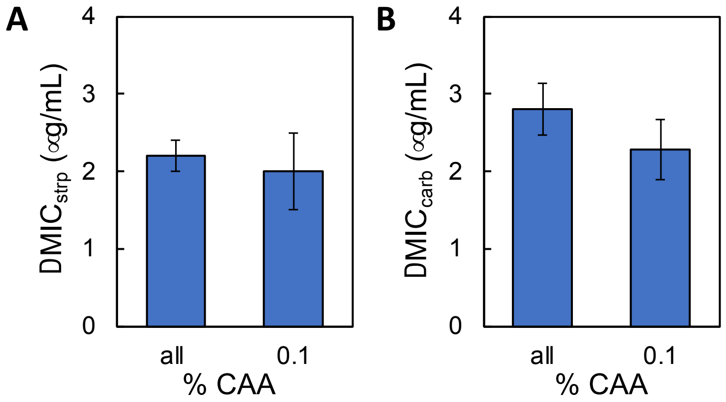


**Supplementary Figure 13: Control data for Figure 6.** ΔMIC of streptomycin (strp, panel A), and carbenicillin (carb, panel B) over the range of casamino acids (% CAA, 0.01%-1%) tested and using only 0.1% casamino acids. For each antibiotic when ΔMIC using all % CAA is compared to ΔMIC using 0.1%, $P \geq 0.38$, two-tailed t-test.
