## Supplementary Fig. 14 for "Purine and pyrimidine synthesis differently affect the strength of the inoculum effect for aminoglycoside and β-lactam antibiotics"

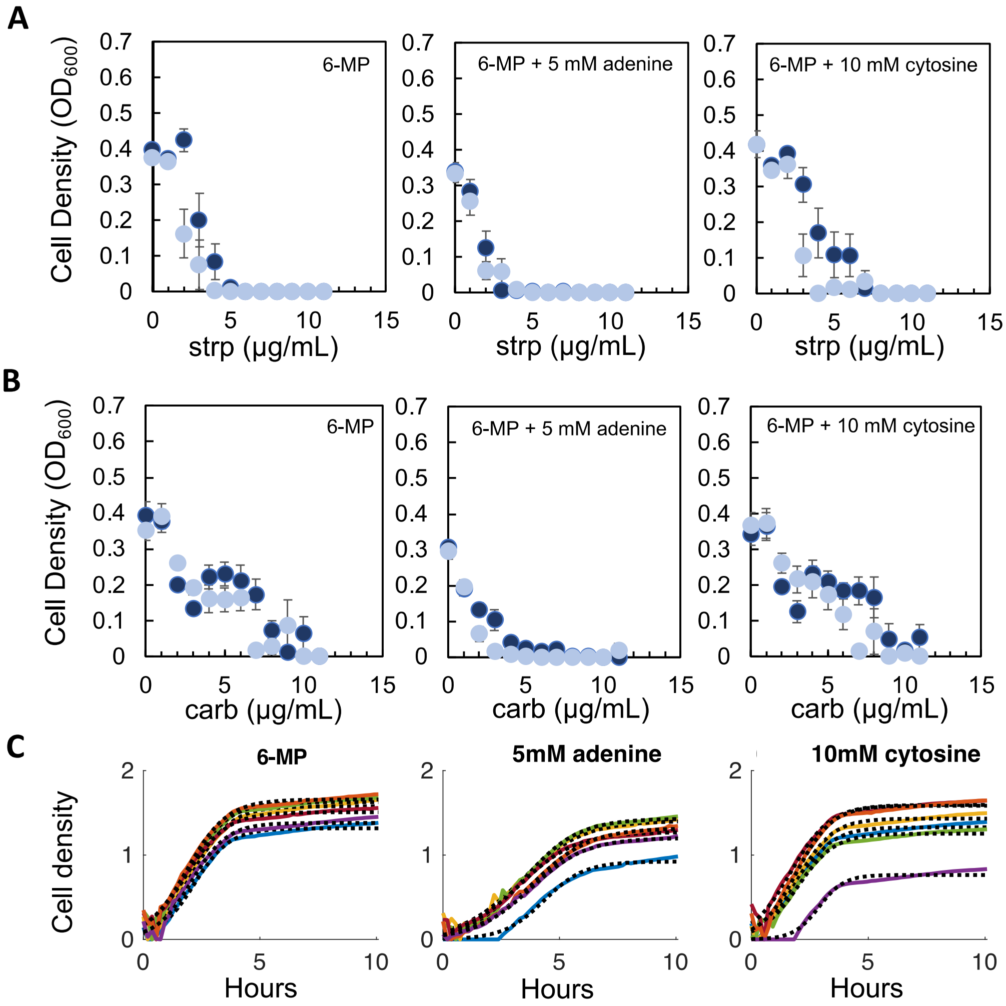


**Supplementary Figure 14: Raw data for experiments using 6-MP.**

**A)** Raw data from which the ΔMIC of streptomycin (strp) was determined when bacteria were challenged with 6-MP. Dark blue = high density, light blue = low density. Error bars = SEM. Average from ≥ 6 biological replicates.

**B)** Raw data from which the ΔMIC of carbenicillin (carb) was determined when bacteria were challenged with 6-MP. Dark blue = high density, light blue = low density. Error bars = SEM. Average from ≥ 7 biological replicates.

**C)** Growth curves of bacteria treated with 6-MP. Colored lines = experimental data. Black dotted lines = growth curve fit using logistic equation.
