## Supplementary Fig. 15 for "Purine and pyrimidine synthesis differently affect the strength of the inoculum effect for aminoglycoside and β-lactam antibiotics"

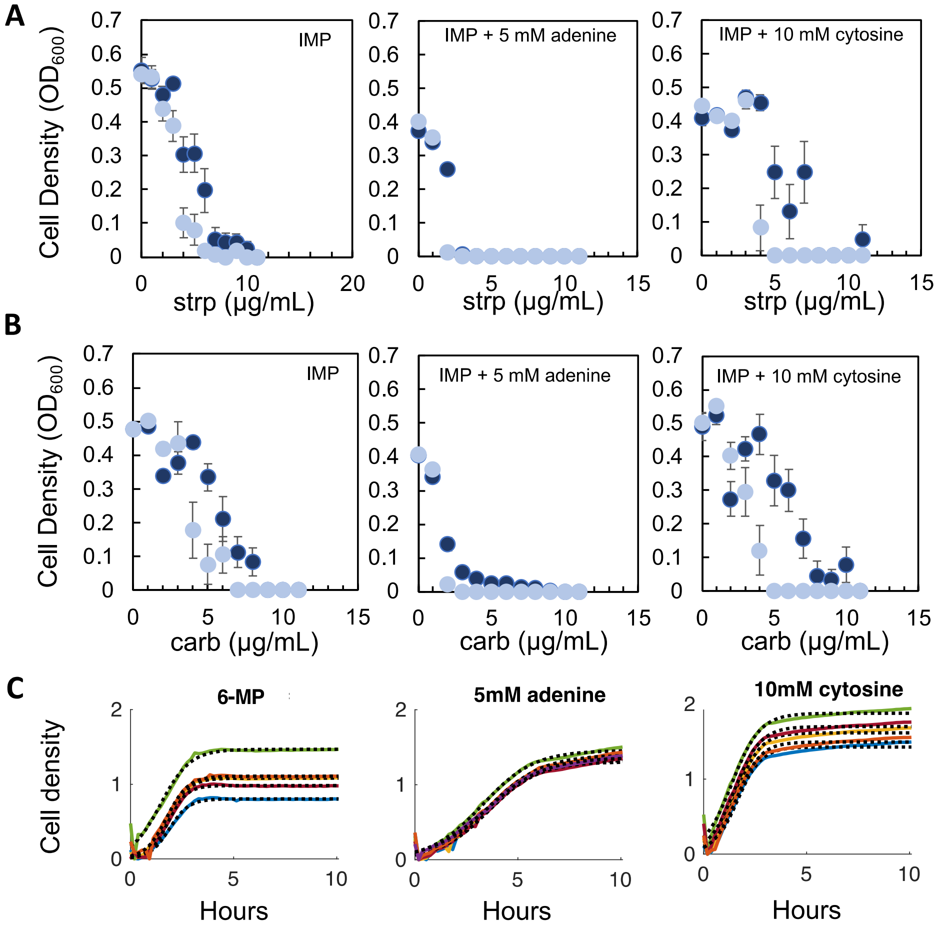


**Supplementary Figure 15: Raw data for experiments using IMP.**

- **A)** Raw data from which the ΔMIC of streptomycin (strp) was determined when bacteria were challenged with IMP. Dark blue = high density, light blue = low density. Error bars = SEM. Average from ≥ 6 biological replicates
- **B)** Raw data from which the ΔMIC of carbenicillin (carb) was determined when bacteria were challenged with IMP. Dark blue = high density, light blue = low density. Error bars = SEM. Average from ≥ 5 biological replicates.
- **C)** Growth curves of bacteria treated with IMP. Colored lines = experimental data. Black dotted lines = growth curve fit using logistic equation.
