## Supplementary Fig. 16 for "Purine and pyrimidine synthesis differently affect the strength of the inoculum effect for aminoglycoside and β-lactam antibiotics"

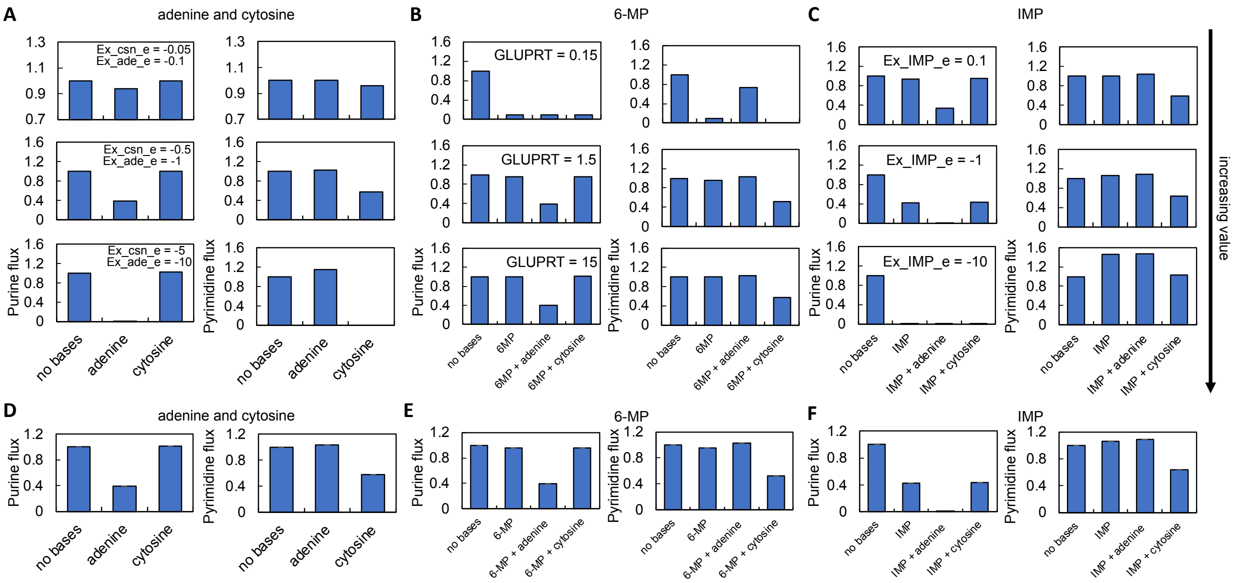


**Supplementary Figure 16: Sensitivity analysis and additional FBA predictions.**

**A)** The effect of changing the lower bound exchange values of cytosine and adenine. Lower bound values are indicated on the plot. For panels A-C, the baseline simulation used for the main text is shown in the middle row. Purine flux represents the reaction catalyzed by PyrC (DHORTS); purine flux represents the reaction catalyzed by PurK (PRAIS).

**B)** The effect of changing the upper bound value of the GLUPRT reaction, which affects the activity of the PurF enzyme. PurF is the target of 6-MP.

**C)** The effect of changing the lower bond value of IMP exchange (Ex_IMP_e).

**D-F)** Average flux values for all reactions involved in nucleotide synthesis up until the synthesis of IMP (purines, after which the pathway branches to produce AMP and GMP) and UMP (pyrimidine). SEM from all reactions. For panels D-F, reactions measured include DHORTS, OMPDC, ORPT, and ASPCT for pyrimidine flux, and IMPC, AIRC3, ADSL2R, AIRC2, GLUPRT, PRAGSR, PRASCSI, and PRAIS for purine flux.
