## Supplementary Fig. 17 for "Purine and pyrimidine synthesis differently affect the strength of the inoculum effect for aminoglycoside and β-lactam antibiotics"

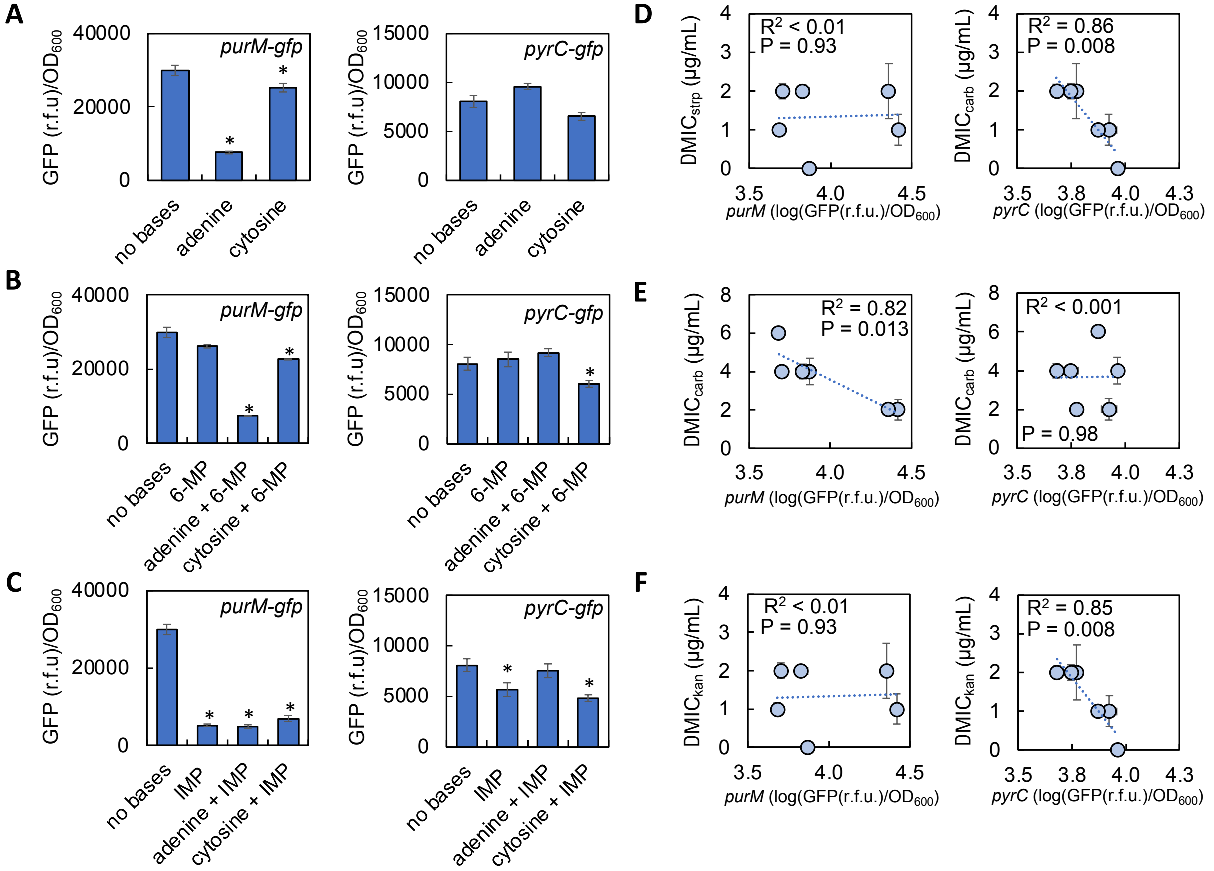


**Supplementary Figure 17: Reporter activity after 24 hours of growth.**

**A)** GFP (in relative fluorescent units, r.f.u.) of *purM*-gfp (left) and *pyrC*-gfp (right) in response to nitrogenous bases. * = different from no nitrogenous base control ($P \leq 0.045$, two-tailed t-test). For all panels, GFP normalized by cell density ($OD_{600}$). Average from 5 biological replicates. Nitrogenous bases that were provided: 5 mM adenine and 10 mM cytosine. Error bars = SEM.

**B)** GFP of *purM*-gfp (left) and *pyrC*-gfp (right) in response to 0.05 µg/mL 6-MP and nitrogenous bases. * = different from no nitrogenous base control ($P \leq 0.041$, two-tailed t-test).

**C)** GFP of *purM*-gfp (left) and *pyrC*-gfp (right) in response to 1 mM IMP and nitrogenous bases. * = different from no nitrogenous base control ($P \leq 0.030$, two-tailed t-test).

**D)** Linear correlation between ΔMIC of streptomycin (strp) and GFP/$OD_{600}$ from either *purM*-gfp (left) and *pyrC*-gfp (right) reporter strains. For panels D-F, P and $R^2$ values from linear regression and ΔMIC from Fig. 6. Weighted least squares (WLS) regression (*purM*: $R^2 < 0.01$, $P = 0.90$; *pyrC*: $R^2 = 0.86$, $P = 0.024$); Deming regression (*purM* - $P = 0.93$; *pyrC* $P = 0.008$).

**E)** Linear correlation between ΔMIC of carbenicillin (carb) and GFP/$OD_{600}$ from either *purM*-gfp (left) and *pyrC*-gfp (right) reporter strains. ΔMIC from Fig. 6. WLS (*purM*: $R^2 = 0.79$, $P = 0.017$; *pyrC*: $R^2 = 0.27$, $P = 0.29$) and Deming regression (*purM* - $P = 0.013$; *pyrC* $P = 0.98$).

F) Linear correlation between ΔMIC of kanamycin (kan) and GFP/$OD_{600}$ from either *purM*-gfp (left) and *pyrC*-gfp (right) reporter strains. ΔMIC from Supplemental Fig. 17. WLS (*purM*: $R^2 < 0.01$, $P = 0.96$; *pyrC*: $R^2 = 0.84$, $P = 0.028$) and Deming regression (*purM* - $P = 0.93$; *pyrC* $P = 0.008$).
