## Supplementary Fig. 18 for "Purine and pyrimidine synthesis differently affect the strength of the inoculum effect for aminoglycoside and β-lactam antibiotics"

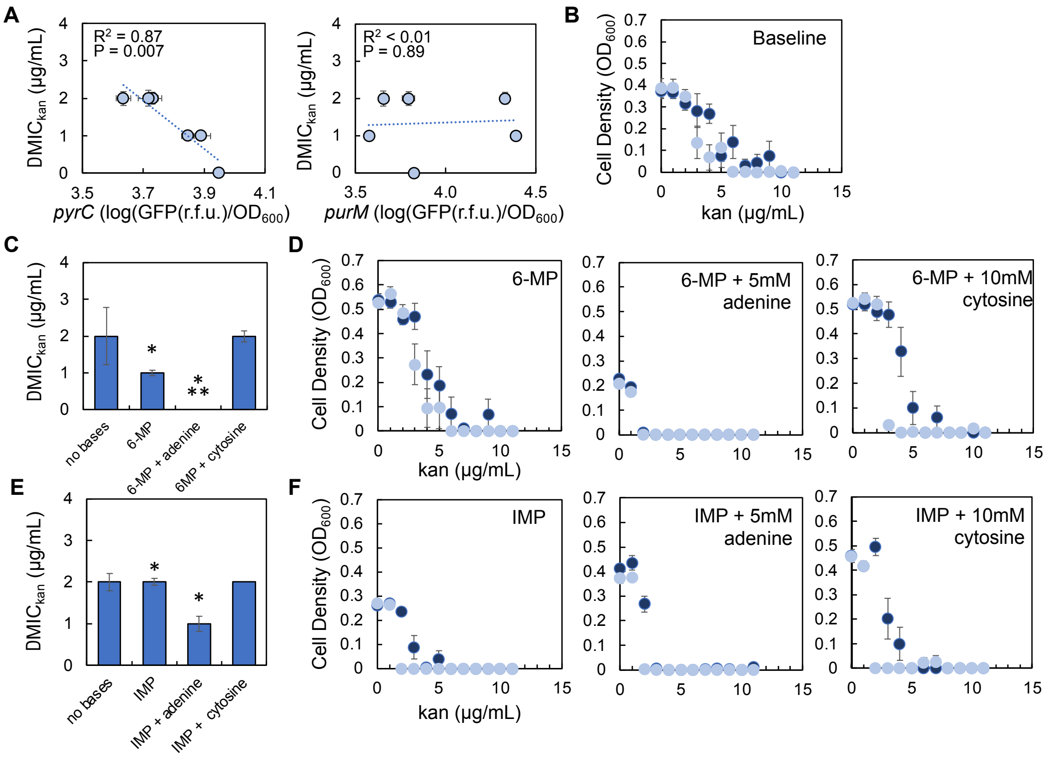


**Supplementary Figure 18: Transcriptional activity of pyrimidine synthesis correlates with ΔMIC of the aminoglycoside kanamycin.**

- **A)** ΔMIC of kanamycin (kan) as a function of *pyrC* (left), and *purM* (right) reporter transcriptional activity. $R^2$ and P value from a linear regression. Weighted least squares (WLS) regression (*purM*: $R^2 < 0.01$, P = 0.99; *pyrC*: $R^2 = 0.87$, P = 0.02); Deming regression (*purM* - P = 0.89; *pyrC* P = 0.007). Error bars = SEM.
- **B)** Raw data from which the baseline ΔMIC of kan was determined. Casamino acids = 0.1%. No nitrogenous bases are included. Dark blue = high density, light blue = low density. Error bars = SEM. Average from = 5 biological replicates.
- **C)** ΔMIC of kan for *E. coli* grown in M9 medium supplemented with 6-MP, 5 mM adenine, and 10 mM cytosine. * = different than no nitrogenous base/no inhibitor control (P ≤ 0.02, two-tailed t-test). ** = not different than zero (P = 0.089, one-tailed t-test). SEM from 5 biological replicates. All P values in Supplementary Table 10.
- **D)** Raw data from which the baseline ΔMIC of kan was determined in the presence of 6-MP. Casamino acids = 0.1%. No nitrogenous bases are included. Dark blue = high density, light blue = low density. Error bars = SEM. Average from = 5 biological replicates.
- **E)** ΔMIC of kan for *E. coli* grown in M9 medium supplemented with IMP, 5 mM adenine, and 10 mM cytosine. * = different than no nitrogenous base/no inhibitor control (P ≤ 0.003, two-tailed t-test). SEM from 6 biological replicates. All P values in Supplementary Table 10.
- **F)** Raw data from which the baseline ΔMIC of kan was determined in the presence of IMP. Casamino acids = 0.1%. No nitrogenous bases are included. Dark blue = high density, light blue = low density. Error bars = SEM. Average from = 6 biological replicates.
