## Supplementary Fig. 19 for "Purine and pyrimidine synthesis differently affect the strength of the inoculum effect for aminoglycoside and β-lactam antibiotics"

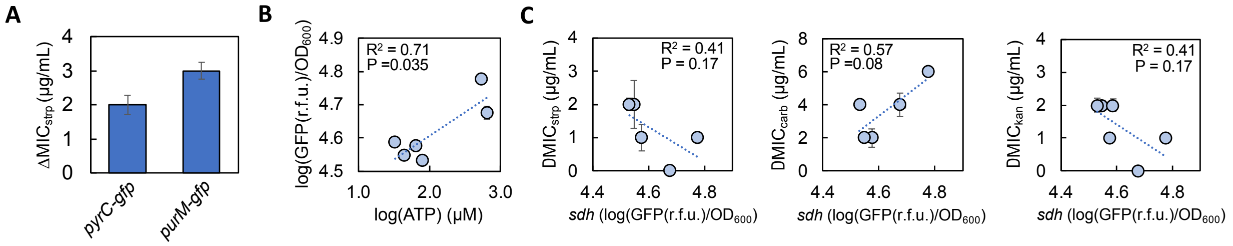


**Supplementary Figure 19: Control experiments and succinate dehydrogenase (*sdh*) promoter activity as determined using reporter strains.**

- **A)** ΔMIC of streptomycin (strp) for *purM-gfp* and *pyrC-gfp* reporter strains. Average from 6 biological replicates. Error bars = SEM.
- **B)** Linear correlation between [ATP] and *sdh*-gfp reporter activity. P and $R^2$ value from linear regression. [ATP] from Fig. 6.
- **C)** Linear correlation between $GFP/OD_{600}$ of *sdh*-gfp reporter strain and ΔMIC of strp, kanamycin (kan), and carbenicillin (carb). ΔMIC for strep and carb from Fig. 3, ΔMIC for kan from Supplementary Fig. 18.
