## Supplementary Methods for "Purine and pyrimidine synthesis differently affect the strength of the inoculum effect for aminoglycoside and β-lactam antibiotics"

*Antibiotics used and their concentrations*

We used the following antibiotics in this study: kanamycin (Fisher Scientific), carbenicillin (Thermofisher), streptomycin (Fisher Scientific), and ciprofloxacin (Thermofisher). For experiments performed with *E. coli*, and except for ciprofloxacin, antibiotics were provided in 1 μg/mL increments. Ciprofloxacin was provided in 0.002 μg/mL increments. For *P. aeruginosa*, streptomycin was provided in 4 μg/mL increments whereas carbenicillin was provided in 5 μg/mL increments.

*pH experiments*

*E. coli* was grown overnight. The following day, the bacteria were washed and inoculated at low initial density into a 96 well plate containing M9 medium with casamino acids (0.01%-1%) and with, or without, nitrogenous bases (5 mM adenine, 10 mM thymine, 10 mM cytosine, 10 mM uracil). We chose to use a low initial density of cells as it allows for the greatest period of growth before reaching stationary. Thus, any changes in pH would be more readily observable that those of a high-density counterpart. After 24 hours of growth (as described in 'MIC assays', main text), 100 μL of the medium was removed, and placed in 5 ml of $dH_2O$ whereupon pH was recorded using a symphony B10P pH meter (VWR, Radnor, PA) that was calibrated right before measurements took place. Similarly, 100 μL of cell-free medium was placed in 5 mL of $dH_2O$, and initial pH was recorded.

*RNA extractions*

*E. coli* was grown overnight whereupon it was diluted 100-fold in 200 μL of fresh M9 medium in a 96 well plate. Two Breathe Easy sealing membranes were placed over top. After 5 hours of growth at 37°C (250RPM), cell density was recorded using $OD_{600}$, and 200 μL of three technical replicates were combined in a 1.5 mL microcentrifuge tube. The cells were centrifuged for 2 minutes at 12,000 RPM and were then resuspended in 30 μL of lysozyme (10 μg/mL, MP Biomedicals). The tube was incubated at room temperature for 20 minutes with vortexing completed every 2 minutes. Total RNA was then extracted using a ZR Fungal/Bacterial RNA MiniPrep kit (Zymo Research) including the optional in-column DNase digest according to the manufacturer's recommendations. Total RNA was quantified using a Qubit fluorometer (ThermoFisher) using the broad range (BR) RNA quantification kit (ThermoFisher) and was normalized by cell density ($OD_{600}$).

*Measuring growth rate*

As performed previously(*1*), $OD_{600}$ values were log-transformed and normalized to the initial minimum density, which removes artifacts and ensures that all growth data were initiated from the same starting point. Together, this helps to reduce the amount of error during curve fitting. Next, we used the logistic equation to determine the maximum growth rate over 10 hours of growth, which allowed most conditions to reach the stationary phase.

$$y = \frac{A}{\left\{1+\exp\left(\frac{4\mu_m}{A}(\lambda-t)+2\right)\right\}} \qquad \text{(Eq. S1)}$$

$$y = A\exp\left\{-\exp\left(\left(\frac{\mu_m e}{A}\right)(\lambda - t) + 1\right)\right\} \qquad \text{(Eq. S2)}$$

where $A$ represents the maximum cell density, $\mu_m$ represents the maximal growth rate, and $\lambda$ represents the lag time. Lower bounds of 0, and upper bounds of 2, 1, and 10 for $A$, $\mu_m$, and $\lambda$ respectively, were used. This step ensures that all parameters are within a biologically feasible parameter space. We then used MATLAB 2023b using the lsqcurvefit function, which is a non-linear least squares solver and serves to estimate values for $A$, $\mu_m$, and $\lambda$ that minimize the differences between the experimental data and fit. After fitting, we determined the average residual. Growth curves with residual values of 0.5 or greater were considered to lack a rigorous fit and were not included in our analysis. Owing to the challenge of fitting both fast (wildtype) and very slow (mutant) growing bacteria using a single model, this was relaxed in the case of the Δ*pyrC* and Δ*purK* mutants such that the mean average residual across all nitrogenous base concentrations was 0.5 or less.
