## Supplementary Results for "Purine and pyrimidine synthesis differently affect the strength of the inoculum effect for aminoglycoside and β-lactam antibiotics"

*Using a 0.01 cutoff for MIC experiments*

After blanking $OD_{600}$ for MIC assays, we set any condition that did not exceed a value of 0.01 to zero. This threshold was used to prevent fluctuations in $OD_{600}$ values observed in cell-free medium counting as growth. Our previous work showed that $OD_{600}$ of cell-free medium could fluctuate +/- 0.009. Thus, values below 0.01 are not robust indicators of bacterial growth. We showed previously that using this cutoff does not impact the general trends in ΔMIC(*1*).

*Nitrogenous base import and downstream effects in E. coli*

Previous research has demonstrated that both purines and pyrimidines provided in the growth medium can be imported into bacteria and affect nucleotide synthesis. For example, *E. coli* grown in minimal medium exogenously supplied with either 1 mM adenine (purine) or 1 mM uracil (pyrimidine) showed cytoplasmic accumulation of both nitrogenous bases using LC-MS/MS(*2*). Adenine can be imported into the cell using PurP(*3*), which can be converted to either AMP or inosine monophosphate (IMP) using purine nitrogenous base salvage pathways, both of which participate in purine biosynthesis. Accumulation of AMP reduces purine synthesis by repressing the activity of PurF. Cytosine can be imported into the cell using CodA/CodB(*4*), and is subsequently converted to uracil via CodA(*5*). Uracil is imported into the cell using UraA(*6*). In both cases, uracil is converted to UMP by Upp. UMP then participates in UTP and CTP *de novo* synthesis. Thymine can also be imported into bacteria using the *rut* pathway; RutG has been shown to import thymine at a rate of 0.3 nmol/mg, which is achieved after ~45 minutes of incubation(*7*). Upon import, thymine is converted to deoxythymidine by DeoA, which is subsequently converted to dTMP by Tdk (*8, 9*). *De novo* pyrimidine synthesis creates UMP, which through a series of reactions leads to the creation of dTMP by Tdk and

ThyA*(9)*. Accumulation of UMP inhibits additional pyrimidine synthesis by inhibiting CarA/CarB whereas the accumulation of dTTP inhibits Tdk activity (*10*). 5-phospho-α-D-ribose 1-diphosphate (*prpp*) serves as a link between *de novo* purine and pyrimidine synthesis as it is used as a substrate for both pathways, thus allowing both pathways to autoregulate (*11*). A reduction in activity in one pathway (e.g., purine synthesis) leads to the accumulation of prpp, which can subsequently be used for the other pathway (e.g., pyrimidine synthesis). Accordingly, and consistent with previous work(*2*), reducing the activity of pyrimidine synthesis through exogenously supplied pyrimidines increases purine synthesis. Similarly, reducing purine synthesis through exogenously supplied purines increases pyrimidine synthesis.

*Alternative explanations for IE*

ATP production: Previous work has shown that when the growth rate is held constant, increasing bacterial metabolism, as measured primarily through ATP production, is a better predictor of antibiotic lethality(*12*). While our previous work showed that growth productivity (Δ[ATP]/Δgrowth rate) is a better predictor of ΔMIC than log[ATP] alone(*1*), we wanted to test the relationship between log[ATP] and ΔMIC in the context of this study where exogenous nitrogenous bases are being provided in the growth medium. While we found significant relationships between log[ATP] and ΔMIC of streptomycin and carbenicillin for *E. coli*, the $R^2$ values (or the strength of the relationship) were lower than that of ΔMIC and log[ATP]/growth rate (Supplementary Fig. 8). We did not find a significant relationship between log[ATP] and ΔMIC of ciprofloxacin. We also did not find a significant relationship between log[ATP] and ΔMIC of streptomycin and carbenicillin for *P. aeruginosa* (Supplementary Fig. 10). Finally, a significant increase in [ATP] absent changes in growth rate relative to the no nitrogenous base control (1 mM adenine, Supplementary Fig. 8) did not alter ΔMIC for streptomycin. Taken together, changes in [ATP] alone cannot fully explain IE.

Growth rate: Previous research has suggested that changes in growth rate alone can alter antibiotic lethality(*13*). While our previous work showed that growth rate alone cannot account for IE and changes in ΔMIC(*1*), we sought to test this in the context of this study. Linear regression analysis indicated that the relationship between growth rate and ΔMIC of streptomycin in *E. coli* was significant but less strong as compared to log[ATP]/growth rate (Supplementary Fig. 8). We did not find a significant relationship between growth rate and ΔMIC of ciprofloxacin. Interestingly, we found a significant relationship between growth rate and ΔMIC of carbenicillin in *E. coli*, the strength of which was greater than that of ΔMIC of log[ATP]/growth rate. This may be owing to the strong dependence of growth rate on the efficacy of β-lactam antibiotics, which has been reported previously(*14*) and deserves future exploration. In addition, we found a moderately significant, but less strong, relationship between growth rate and ΔMIC of streptomycin for *P. aeruginosa* (Supplementary Fig. 10). The same was not found when *P. aeruginosa* was challenged with carbenicillin. Finally, a significant reduction in growth rate absent changes in [ATP] relative to the no nitrogenous base control (0.9 mM thymine, Supplementary Fig. 8) did not alter ΔMIC for streptomycin. Taken together, growth rate is not the most consistent predictor of ΔMIC when considering multiple bacterial species and antibiotics.

Carrying capacity: As the production of ATP is growth phase-dependent(*15, 16*), a higher carrying capacity in the growth medium may allow an extended log phase, which could impact antibiotic lethality and ΔMIC. To test the relationship between the carrying capacity of the growth medium (determined using the average $OD_{600}$ reached by the high and low initial density populations in antibiotic-free medium), we performed a linear regression between ΔMIC and carrying capacity. For *E. coli*, we found significant relationships between carrying capacity and ΔMIC of streptomycin and carbenicillin (Supplementary Fig. 9). However, the strength of these relationships was less than the relationships between ΔMIC and log[ATP]/growth rate. We did not find a significant relationship between carrying capacity and ΔMIC of ciprofloxacin. Taken together, carrying capacity alone is unlikely to be the strongest determinant of IE.

pH of the growth medium: Previous studies have shown that pH influences bacterial metabolism (*17*) and antibiotic lethality(*18*). As we did not use a buffered medium in our analysis, we wanted to ensure that alterations to pH due to bacterial growth or differences in media composition were not driving our results. Accordingly, we measured the pH of the medium both before and after bacterial growth (see Supplementary Methods). The medium that was tested contained no nitrogenous bases, 5 mM adenine, 10 mM cytosine, 10 mM thymine, or 10 mM uracil. We averaged pH across the five percentages of casamino acids (0.01, 0.05, 0.1, 0.5, and 1%) and plotted the values against ΔMIC of streptomycin, carbenicillin, and ciprofloxacin. We did not find a significant nor strong relationship between ΔMIC and either initial or final pH (Supplementary Fig. 9), which was consistent with our previous work(*1*).

Cell-cell communication: Previous work has indicated that secreted products produced by bacteria can confer antibiotic tolerance and resistance. These include molecules that create persister cells(*19*) and small molecules produced during quorum sensing(*20*). We previously showed that the removal of *tnaA*, which is implicated in the production of indole and the formation of persister cells(*21*), does not abolish IE(*22*). To test the effect of removing quorum sensing, we used a knockout strain that lacks *luxS*, which is involved in AI-2 production in *E. coli* (*23*). We found that when grown without nitrogenous bases or with 10 mM cytosine, IE continued to be present (Supplementary Fig. 9). When grown with 10 mM thymine, and consistent with the wildtype strain containing *luxS*, ΔMIC was reduced relative to the no nitrogenous base control. Thus, quorum sensing does not appear to account for IE.

Antibiotic target to antibiotic ratio: Previous work has suggested that the ratio of antibiotic to antibiotic target can account for IE(*24*). For a given concentration of antibiotic, the greater the density of the population, the fewer antibiotic molecules per antibiotic target, which leads to increased tolerance. While our previous work provided evidence that this alternative explanation could not account for IE(*1, 22*), we explicitly tested this hypothesis in the context of our study by quantifying the total amount of RNA extracted from bacteria. Total RNA can be used as a reliable surrogate of the quantity of rRNA in the cell as 95–97% of total RNA is rRNA in bacteria(*25*) and tRNAs are removed during the purification process. We extracted total RNA from *E. coli* grown without exogenous nitrogenous bases, with 10 mM cytosine and with 10 mM thymine. We then correlated the quantity of total RNA normalized by cell density to ΔMIC of streptomycin; note that the molecular target of streptomycin is the ribosome. We did not find a significant nor strong relationship between the quantity of total RNA and ΔMIC of streptomycin (Supplementary Fig. 9). This indicates that changes in the number of molecular targets as a result of exogenously applied nitrogenous bases cannot explain our findings.

Efflux pumps: Recent work has shown that the presence of efflux pumps can protect neighboring bacteria from the effects of antibiotics(*26*). Thus, it is possible that efflux pumps could account for density-dependent antibiotic resistance; conceivably, the higher the population density, the more efflux pumps would be available to collectively protect the population. To test this, we acquired a knockout strain that lacks the *mdtA* efflux pump, which confers antibiotic resistance(*27*). ΔMIC of this mutant was consistent when 10 mM cytosine was included in the medium as compared to a no nitrogenous base control (Supplementary Fig. 9). Supplementation with 10 mM thymine reduced ΔMIC of the knockout strain as compared to the control. Our previous work has also shown that deletion of *tolC*, *acrA,* and *acrB*, all of which encode efflux pumps, continues to allow IE. Thus, efflux pumps are unlikely the leading cause of IE in our system(*1*).

<u>*ampC* β-lactamase</u>: *E. coli* strain BW25113 contains an endogenous *ampC* β-lactamase. However, carbenicillin is not readily broken down by the *ampC* β-lactamase (*28*). Moreover, our previous work showed that without the *ampC* β-lactamase (Δ*ampC*), IE continues to be present for carbenicillin in BW25113(*1*). Taken together, AmpC is not playing a large role in determining IE for carbenicillin.

*Flux balance analysis – parameter estimation*

We performed both flux balance analysis and flux balance analysis with Optknock, using the COBRA toolbox v.3.0(*29*) and the iML1515 model of *E. coli* (*30*). All lower and upper bound values that are considered standard in the iML1515 model were not altered. We altered the lower bound exchange values of $K^+$, $Mg^{2+}$, $Na^+$, $NH_4^+$, $Cl^-$ $P_i$, $SO_4^{2-}$, and $Ca^{2+}$ to -1000 to account for the composition of M9 medium.

All lower bound exchange values were determined from previously published work and can be found in Supplementary Table 1. All upper bound values remained 1000 except in the case where 6-MP was used in the medium (see *Flux balance analysis – simulations*). When determining $g_{DW}$ from previous work, we assumed that 1 $OD_{600}$ is approximately 0.39g/L cell dry weight (*31*). The lower bound exchange flux for amino acids was estimated as described previously(*1*) using published data(*32*). We approximated the order of magnitude of the lower bound flux value of glucose using previously published work (*33, 34*). The oxygen consumption rate (O2_tex) was estimated from previously published work(*1, 35*). Finally, the thiamine consumption rate was determined from previous literature (*36*).

To estimate the lower bound flux values of nitrogenous bases provided in the growth medium, we used previously published work to estimate the order of magnitude of uptake when ~ 1 μM of each nitrogenous base was separately provided to *E. coli* (*3, 37-40*). To account for experiments where the concentration of nitrogenous bases provided in the medium was 1-10 mM, we considered previous data showing that increasing the concentration of nitrogenous bases increases uptake near linearly(*39*), but can also saturate once higher concentrations (~0.8 mM) are reached(*3*). To take into account both trends, we first scaled the order of magnitude of the lower bound flux value linearly as a function of increasing concentration of nitrogenous bases. However, to account for the saturation of uptake at high concentrations of nitrogenous bases, we decreased the lower bound value one order of magnitude from the linear scaling.

*Flux balance analysis – simulations*

We performed both flux balance analysis and flux balance analysis with Optknock, using the COBRA toolbox v.3.0 (*29*) and the iML1515 model of *E. coli* (*30*). MATLAB 2019b was used for simulations owing to its established compatibility with multiple optimization solvers. The LP solver of Gurobi was used as the optimization software (version 11.0.0). We used git version 2.39.2. For all simulations, biomass was set as the objective function.

For Optknock simulations, all 1506 genes were removed, and resulting changes in ATPsyn and biomass were evaluated. Biomass values of 0 were set to $10^{-14}$, which was less than the lowest biomass value simulated ($-1.26 \times 10^{-13}$). This was performed so that ATPsyn/biomass could be determined for genes whose removal was lethal (biomass = 0). Pathways, as identified using Ecocyc,(*41*) were then assigned to genes whose removal increased ATPsyn/biomass above that of the wildtype (simulations performed without knocking out any genes but using the same parameter set). We then calculated the frequency of the occurrence of each pathway and their average ATP/syn values

(Fig. 1). Importantly, this was performed only for genes whose removal increased ATPsyn/biomass above wildtype.

For simulations that examined flux through purine and pyrimidine synthesis, we measured the flux of DHORTS and PRAIS (Fig. 7A). The DHORTS reaction captures the activity of the PyrC dihydroorotase. The PRAIS reaction captures the activity of the PurM phosphoribosylaminoimidazole synthase. Together, these reactions match the promoters of genes that were used for the reporter strain assays (*pyrC* and *purM* promoters). We note that simulation predictions with DHORTS and PRAIS are consistent when multiple reactions in pyrimidine (DHORTS, OMPDC, ORPT, AND ASPCT) and purine (IMPC, AIRC3, ADSL2R, AIRC2, GLUPRT, PRAGSR, PRASCSI, and PRAIS) synthesis are considered (Supplementary Fig. 18).

For simulations with 6-MP (Fig. 7), we reduced the upper bound flux value of the GLUPRT reaction (captures the *purF* glutamine phosphoribosyldiphosphate amidotransferase reaction, which is inhibited by 6-MP(*42*)), which approximates the effect of an inhibitor by reducing the maximum flux through this part of purine synthesis. For simulations with IMP (Fig. 7), we modified the lower bound flux value of IMP exchange (EX_imp_e). As IMP uptake rates are not yet reported, we estimated the order of magnitude of this lower bound flux value using previously reported flux values of multiple carbon sources provided to *E. coli* (*1*). Sensitivity analysis for these parameters can be found in Supplementary Fig. 18.
