## Supplementary Table 1 for "Purine and pyrimidine synthesis differently affect the strength of the inoculum effect for aminoglycoside and β-lactam antibiotics"

### Supplementary Tables

**Supplementary Table 1: Upper and lower bound flux values used for FBA simulations in Figs. 1, 6, and 7.** All other flux values were left unchanged in the model.

| Reaction | Lower bound | Upper bound | Ref |
|---|---|---|---|
| **Core reactions** | | | |
| Glycine | -0.8835861 | 1000 | (*32*) |
| Alanine | -1.682609333 | 1000 | (*32*) |
| Arginine | -1.6722119 | 1000 | (*32*) |
| Asparagine | -2.396529233 | 1000 | (*32*) |
| Aspartate | 0.895297803 | 1000 | (*32*) |
| Cysteine | -1.4220841 | 1000 | (*32*) |
| Glutamate | -1.5974812 | 1000 | (*32*) |
| Glutamine | -1.987637733 | 1000 | (*32*) |
| Histidine | -3.3186743 | 1000 | (*32*) |
| Isoleucine | -2.090270167 | 1000 | (*32*) |
| Leucine | -2.090270167 | 1000 | (*32*) |
| Lysine | -2.177075267 | 1000 | (*32*) |
| Methionine | -2.431328733 | 1000 | (*32*) |
| Phenylalanine | -2.422301767 | 1000 | (*32*) |
| Proline | -2.3791969 | 1000 | (*32*) |
| Serine | -0.403842533 | 1000 | (*32*) |
| Threonine | -1.907954967 | 1000 | (*32*) |
| Tryptophan | -3.014569033 | 1000 | (*32*) |
| Tyrosine | -3.524060967 | 1000 | (*32*) |
| Valine | -2.266623633 | 1000 | (*32*) |
| Thiamine | -0.000001 | 1000 | (*36*) |
| Oxygen (O2tex) | -20 | 1000 | (*1, 35*) |
| Glucose | -18 | 1000 | (*33, 34*) |
| **Values specific to Fig. 6 and 7** | | | |
| 6-MP (GLUPRT) | 0 | 0.15 | Supplementary results |
| IMP (Ex_IMP_e) | -1 | 1000 | Supplementary results |
| **Values specific for Fig. 7** | | | |
| Adenine (Ex_ade_e) | -1 | 1000 | Supplementary results |
| Cytosine (Ex_csn_e) | -0.5 | 1000 | Supplementary results |
