## Supplementary Table 2 for "Purine and pyrimidine synthesis differently affect the strength of the inoculum effect for aminoglycoside and β-lactam antibiotics"

**Supplementary Table 2: P values and the exact number of biological replicates (*n*) for Fig. 2.** P value was determined using a two-tailed t-test against the no nitrogenous base control.

| Condition | [nitrogenous base] mM | % CAA | *n* (growth rate) | P value (growth rate) | *n* ([ATP]) | P value ([ATP]) |
|---|---|---|---|---|---|---|
| **No nitrogenous bases** | 0 | 0.01 | 7 | 1 | 4 | 1 |
| | | 0.05 | 8 | | | |
| | | 0.1 | 8 | | | |
| | | 0.5 | 8 | | | |
| | | 1 | 8 | | | |
| **Adenine** | 1 | 0.01 | 7 | 0.567 | | 6.252E-11 |
| | | 0.05 | 7 | | | |
| | | 0.1 | 5 | | | |
| | | 0.5 | 8 | | | |
| | | 1 | 7 | | | |
| | 5 | 0.01 | 8 | 5.47E-16 | | 2.17E-14 |
| | | 0.05 | 8 | | | |
| | | 0.1 | 6 | | | |
| | | 0.5 | 6 | | | |
| | | 1 | 7 | | | |
| | 10 | 0.01 | 7 | 2.30E-21 | | 6.91E-11 |
| | | 0.05 | 5 | | | |
| | | 0.1 | 7 | | | |
| | | 0.5 | 6 | | | |
| | | 1 | 5 | | | |
| **Cytosine** | 1 | 0.01 | 7 | 2.75E-04 | | 8.11E-04 |
| | | 0.05 | 6 | | | |
| | | 0.1 | 6 | | | |
| | | 0.5 | 8 | | | |
| | | 1 | 8 | | | |
| | 5 | 0.01 | 8 | 0.107 | | 0.003 |
| | | 0.05 | 7 | | | |
| | | 0.1 | 7 | | | |
| | | 0.5 | 6 | | | |
| | | 1 | 8 | | | |
| | 10 | 0.01 | 7 | 0.437 | | 0.244 |
| | | 0.05 | 6 | | | |
| | | 0.1 | 8 | | | |
| | | 0.5 | 7 | | | |
| | | 1 | 6 | | | |
| **Thymine** | 1 | 0.01 | 8 | 0.004 | | 4.948E-04 |
| | | 0.05 | 6 | | | |
| | | 0.1 | 8 | | | |
| | | 0.5 | 8 | | | |
| | | 1 | 8 | | | |
| | 5 | 0.01 | 8 | 3.75E-13 | | 1.78E-09 |

| | | | | | | |
|---|---|---|---|---|---|---|
| | | 0.05 | 8 | | | |
| | | 0.1 | 8 | | | |
| | | 0.5 | 7 | | | |
| | | 1 | 7 | | | |
| | 10 | 0.01 | 8 | 1.101E-17 | | 3.644E-11 |
| | | 0.05 | 4 | | | |
| | | 0.1 | 8 | | | |
| | | 0.5 | 8 | | | |
| | | 1 | 5 | | | |
| **Uracil** | 1 | 0.01 | 7 | 0.386 | | 0.002 |
| | | 0.05 | 9 | | | |
| | | 0.1 | 10 | | | |
| | | 0.5 | 10 | | | |
| | | 1 | 12 | | | |
| | 5 | 0.01 | 9 | 0.276 | | 0.002 |
| | | 0.05 | 12 | | | |
| | | 0.1 | 9 | | | |
| | | 0.5 | 8 | | | |
| | | 1 | 12 | | | |
| | 10 | 0.01 | 8 | 0.639 | | 0.004 |
| | | 0.05 | 7 | | | |
| | | 0.1 | 7 | | | |
| | | 0.5 | 11 | | | |
| | | 1 | 12 | | | |
