## Supplementary Table 3 for "Purine and pyrimidine synthesis differently affect the strength of the inoculum effect for aminoglycoside and β-lactam antibiotics"

**Supplementary Table 3: Average residual values for growth curve fitting to determine growth rates in Fig. 2.**

| Growth condition | % CAA | Biological replicate | | | | | | | | | | | | Average | SEM |
|---|---|---|---|---|---|---|---|---|---|---|---|---|---|---|---|
| no nitrogenous bases | **0.01** | 0.22 | 0.09 | 0.04 | 0.02 | 0.05 | 0.09 | 0.06 | - | - | - | - | - | 0.09 | 0.02 |
| | **0.05** | 0.15 | 0.14 | 0.03 | 0.20 | 0.09 | 0.08 | 0.10 | 0.05 | - | - | - | - | 0.13 | 0.02 |
| | **0.1** | 0.16 | 0.20 | 0.26 | 0.18 | 0.11 | 0.12 | 0.15 | 0.14 | - | - | - | - | 0.20 | 0.02 |
| | **0.5** | 0.13 | 0.10 | 0.07 | 0.12 | 0.08 | 0.08 | 0.11 | 0.10 | - | - | - | - | 0.11 | 0.01 |
| | **1** | 0.07 | 0.07 | 0.15 | 0.09 | 0.07 | 0.04 | 0.05 | 0.05 | - | - | - | - | 0.09 | 0.01 |
| 1 mM adenine | **0.01** | 0.03 | 0.04 | 0.18 | 0.04 | 0.06 | 0.14 | 0.25 | - | - | - | - | - | 0.07 | 0.03 |
| | **0.05** | 0.02 | 0.01 | 0.08 | 0.08 | 0.31 | 0.21 | 0.10 | - | - | - | - | - | 0.05 | 0.04 |
| | **0.1** | 0.11 | 0.13 | 0.18 | 0.11 | 0.45 | - | - | - | - | - | - | - | 0.13 | 0.06 |
| | **0.5** | 0.08 | 0.06 | 0.06 | 0.07 | 0.09 | 0.08 | 0.20 | 0.27 | - | - | - | - | 0.07 | 0.03 |
| | **1** | 0.08 | 0.04 | 0.06 | 0.09 | 0.13 | 0.11 | 0.08 | - | - | - | - | - | 0.07 | 0.01 |
| 5 mM adenine | **0.01** | 0.33 | 0.11 | 0.09 | 0.08 | 0.02 | 0.03 | 0.18 | 0.10 | - | - | - | - | 0.15 | 0.03 |
| | **0.05** | 0.01 | 0.08 | 0.14 | 0.09 | 0.12 | 0.18 | 0.22 | - | - | - | - | - | 0.08 | 0.02 |
| | **0.1** | 0.33 | 0.08 | 0.06 | 0.05 | 0.05 | 0.02 | - | - | - | - | - | - | 0.13 | 0.04 |
| | **0.5** | 0.13 | 0.19 | 0.15 | 0.04 | 0.08 | 0.08 | - | - | - | - | - | - | 0.13 | 0.02 |
| | **1** | 0.16 | 0.17 | 0.05 | 0.08 | 0.06 | 0.08 | 0.05 | - | - | - | - | - | 0.11 | 0.02 |
| 10 mM adenine | **0.01** | 0.02 | 0.07 | 0.19 | 0.05 | 0.16 | 0.11 | 0.12 | - | - | - | - | - | 0.08 | 0.02 |
| | **0.05** | 0.25 | 0.27 | 0.02 | 0.07 | 0.14 | - | - | - | - | - | - | - | 0.15 | 0.04 |
| | **0.1** | 0.15 | 0.15 | 0.02 | 0.12 | 0.08 | 0.18 | 0.01 | - | - | - | - | - | 0.11 | 0.02 |
| | **0.5** | 0.04 | 0.13 | 0.17 | 0.14 | 0.26 | 0.13 | - | - | - | - | - | - | 0.12 | 0.03 |
| | **1** | 0.16 | 0.01 | 0.10 | 0.04 | 0.05 | - | - | - | - | - | - | - | 0.08 | 0.02 |
| 1 mM cytosine | **0.01** | 0.05 | 0.05 | 0.03 | 0.16 | 0.03 | 0.13 | 0.13 | - | - | - | - | - | 0.07 | 0.02 |
| | **0.05** | 0.07 | 0.07 | 0.04 | 0.28 | 0.25 | 0.17 | - | - | - | - | - | - | 0.11 | 0.04 |
| | **0.1** | 0.20 | 0.25 | 0.06 | 0.30 | 0.14 | 0.27 | - | - | - | - | - | - | 0.20 | 0.03 |
| | **0.5** | 0.08 | 0.11 | 0.07 | 0.10 | 0.07 | 0.09 | 0.15 | 0.10 | - | - | - | - | 0.09 | 0.01 |
| | **1** | 0.07 | 0.06 | 0.05 | 0.24 | 0.11 | 0.08 | 0.09 | 0.06 | - | - | - | - | 0.11 | 0.02 |
| 5 mM cytosine | **0.01** | 0.05 | 0.09 | 0.04 | 0.05 | 0.02 | 0.03 | 0.12 | 0.14 | - | - | - | - | 0.06 | 0.01 |
| | **0.05** | 0.10 | 0.12 | 0.06 | 0.36 | 0.12 | 0.21 | 0.06 | - | - | - | - | - | 0.16 | 0.04 |
| | **0.1** | 0.09 | 0.08 | 0.22 | 0.08 | 0.15 | 0.22 | 0.13 | - | - | - | - | - | 0.12 | 0.02 |
| | **0.5** | 0.09 | 0.09 | 0.10 | 0.08 | 0.24 | 0.41 | - | - | - | - | - | - | 0.09 | 0.05 |
| | **1** | 0.03 | 0.15 | 0.11 | 0.35 | 0.13 | 0.23 | 0.16 | 0.08 | - | - | - | - | 0.16 | 0.03 |
| 10 mM cytosine | **0.01** | 0.16 | 0.11 | 0.02 | 0.07 | 0.04 | 0.08 | 0.06 | - | - | - | - | - | 0.09 | 0.02 |
| | **0.05** | 0.25 | 0.09 | 0.11 | 0.08 | 0.22 | 0.12 | - | - | - | - | - | - | 0.13 | 0.03 |
| | **0.1** | 0.20 | 0.25 | 0.12 | 0.12 | 0.17 | 0.09 | 0.12 | 0.09 | - | - | - | - | 0.17 | 0.02 |
| | **0.5** | 0.21 | 0.12 | 0.32 | 0.12 | 0.09 | 0.12 | 0.09 | - | - | - | - | - | 0.19 | 0.03 |
| | **1** | 0.12 | 0.12 | 0.09 | 0.08 | 0.08 | 0.05 | - | - | - | - | - | - | 0.10 | 0.01 |
| 1 mM thymine | **0.01** | 0.04 | 0.11 | 0.19 | 0.15 | 0.05 | 0.06 | 0.12 | 0.15 | - | - | - | - | 0.12 | 0.02 |
| | **0.05** | 0.12 | 0.25 | 0.12 | 0.23 | 0.17 | 0.11 | - | - | - | - | - | - | 0.18 | 0.02 |

| | | | | | | | | | | | | | | |
|---|---|---|---|---|---|---|---|---|---|---|---|---|---|---|
| | **0.1** | 0.24 | 0.17 | 0.13 | 0.10 | 0.20 | 0.13 | 0.23 | 0.21 | - | - | - | - | 0.16 | 0.02 |
| | **0.5** | 0.11 | 0.21 | 0.10 | 0.12 | 0.07 | 0.23 | 0.16 | 0.10 | - | - | - | - | 0.14 | 0.02 |
| | **1** | 0.05 | 0.28 | 0.06 | 0.05 | 0.25 | 0.13 | 0.16 | 0.08 | - | - | - | - | 0.11 | 0.03 |
| | **0.01** | 0.09 | 0.15 | 0.09 | 0.20 | 0.04 | 0.08 | 0.13 | 0.09 | - | - | - | - | 0.13 | 0.02 |
| | **0.05** | 0.23 | 0.32 | 0.14 | 0.11 | 0.14 | 0.01 | 0.02 | 0.13 | - | - | - | - | 0.20 | 0.03 |
| | **0.1** | 0.19 | 0.16 | 0.01 | 0.30 | 0.00 | 0.16 | 0.15 | 0.00 | - | - | - | - | 0.16 | 0.04 |
| **5 mM thymine** | **0.5** | 0.09 | 0.14 | 0.11 | 0.27 | 0.12 | 0.18 | 0.09 | - | - | - | - | - | 0.15 | 0.02 |
| | **1** | 0.14 | 0.09 | 0.06 | 0.06 | 0.13 | 0.18 | 0.20 | - | - | - | - | - | 0.09 | 0.02 |
| | **0.01** | 0.24 | 0.08 | 0.04 | 0.06 | 0.05 | 0.05 | 0.08 | 0.06 | - | - | - | - | 0.11 | 0.02 |
| | **0.05** | 0.13 | 0.10 | 0.15 | 0.09 | - | - | - | - | - | - | - | - | 0.12 | 0.01 |
| | **0.1** | 0.30 | 0.35 | 0.10 | 0.07 | 0.13 | 0.11 | 0.15 | 0.10 | - | - | - | - | 0.21 | 0.03 |
| **10 mM thymine** | **0.5** | 0.25 | 0.21 | 0.12 | 0.07 | 0.07 | 0.14 | 0.21 | 0.21 | - | - | - | - | 0.16 | 0.02 |
| | **1** | 0.11 | 0.17 | 0.11 | 0.13 | 0.08 | - | - | - | - | - | - | - | 0.13 | 0.01 |
| | **0.01** | 0.01 | 0.00 | 0.03 | 0.07 | 0.01 | 0.01 | 0.01 | - | - | - | - | - | 0.03 | 0.01 |
| | **0.05** | 0.05 | 0.09 | 0.06 | 0.25 | 0.22 | 0.10 | 0.03 | 0.03 | 0.36 | - | - | - | 0.11 | 0.04 |
| | **0.1** | 0.07 | 0.07 | 0.09 | 0.06 | 0.11 | 0.12 | 0.38 | 0.34 | 0.37 | 0.30 | - | - | 0.07 | 0.04 |
| | **0.5** | 0.08 | 0.10 | 0.33 | 0.05 | 0.21 | 0.17 | 0.11 | 0.05 | 0.07 | 0.07 | - | - | 0.14 | 0.03 |
| **1 mM uracil** | **1** | 0.06 | 0.11 | 0.11 | 0.09 | 0.12 | 0.10 | 0.16 | 0.14 | 0.04 | 0.05 | 0.05 | 0.04 | 0.09 | 0.01 |
| | **0.01** | 0.03 | 0.47 | 0.04 | 0.01 | 0.14 | 0.00 | 0.03 | 0.01 | 0.01 | - | - | - | 0.14 | 0.05 |
| | **0.05** | 0.04 | 0.26 | 0.17 | 0.35 | 0.18 | 0.13 | 0.21 | 0.12 | 0.09 | 0.25 | 0.32 | 0.35 | 0.21 | 0.03 |
| | **0.1** | 0.27 | 0.14 | 0.08 | 0.08 | 0.09 | 0.24 | 0.24 | 0.20 | 0.24 | - | - | - | 0.14 | 0.02 |
| **5 mM uracil** | **0.5** | 0.06 | 0.13 | 0.12 | 0.10 | 0.10 | 0.07 | 0.07 | 0.07 | - | - | - | - | 0.10 | 0.01 |
| | **1** | 0.07 | 0.07 | 0.09 | 0.07 | 0.14 | 0.07 | 0.08 | 0.08 | 0.05 | 0.04 | 0.05 | 0.05 | 0.07 | 0.01 |
| | **0.01** | 0.04 | 0.05 | 0.02 | 0.15 | 0.01 | 0.06 | 0.04 | 0.13 | - | - | - | - | 0.06 | 0.02 |
| | **0.05** | 0.04 | 0.13 | 0.29 | 0.05 | 0.04 | 0.37 | 0.05 | - | - | - | - | - | 0.13 | 0.05 |
| | **0.1** | 0.04 | 0.36 | 0.17 | 0.20 | 0.08 | 0.06 | 0.41 | - | - | - | - | - | 0.19 | 0.05 |
| **10 mM uracil** | **0.5** | 0.05 | 0.09 | 0.11 | 0.13 | 0.31 | 0.15 | 0.36 | 0.10 | 0.08 | 0.09 | 0.09 | - | 0.10 | 0.03 |
| | **1** | 0.13 | 0.07 | 0.07 | 0.10 | 0.07 | 0.05 | 0.05 | 0.05 | 0.06 | 0.04 | 0.05 | 0.05 | 0.09 | 0.01 |
