## Supplementary Table 4 for "Purine and pyrimidine synthesis differently affect the strength of the inoculum effect for aminoglycoside and β-lactam antibiotics"

**Supplementary Table 4: Average residual values for growth curve fitting to determine growth rates for *P. aeruginosa* in Fig. 4.**

| Growth condition | % CAA | Biological replicate | | | | | | Average | SEM |
|---|---|---|---|---|---|---|---|---|---|
| no nitrogenous bases | | | | | | | | | |
| | **0.1** | 0.26 | 0.15 | 0.33 | 0.41 | 0.15 | - | 0.26 | 0.04 |
| | **0.5** | 0.27 | 0.32 | 0.49 | 0.22 | 0.25 | 0.46 | 0.34 | 0.04 |
| | **1** | 0.17 | 0.22 | 0.45 | 0.11 | 0.27 | 0.17 | 0.23 | 0.04 |
| 5 mM adenine | | | | | | | | | |
| | **0.1** | 0.27 | 0.16 | 0.00 | 0.17 | 0.29 | - | 0.18 | 0.05 |
| | **0.5** | 0.16 | 0.24 | 0.42 | 0.15 | 0.30 | 0.16 | 0.24 | 0.04 |
| | **1** | 0.22 | 0.23 | 0.41 | 0.22 | 0.36 | 0.32 | 0.29 | 0.03 |
| 10 mM cytosine | | | | | | | | | |
| | **0.1** | 0.18 | 0.27 | 0.22 | 0.27 | 0.19 | - | 0.23 | 0.02 |
| | **0.5** | 0.20 | 0.35 | 0.13 | 0.19 | 0.21 | - | 0.21 | 0.03 |
| | **1** | 0.41 | 0.28 | 0.21 | 0.31 | 0.21 | - | 0.28 | 0.03 |
| 10 mM thymine | | | | | | | | | |
| | **0.1** | 0.15 | 0.13 | 0.22 | 0.25 | 0.23 | - | 0.19 | 0.02 |
| | **0.5** | 0.12 | 0.37 | 0.17 | 0.12 | - | - | 0.20 | 0.05 |
| | **1** | 0.21 | 0.29 | 0.43 | 0.20 | 0.30 | 0.40 | 0.31 | 0.04 |
| 10 mM uracil | | | | | | | | | |
| | **0.1** | 0.16 | 0.21 | 0.20 | 0.45 | 0.23 | - | 0.25 | 0.05 |
| | **0.5** | 0.11 | 0.13 | 0.33 | 0.20 | 0.19 | 0.19 | 0.19 | 0.03 |
| | **1** | 0.14 | 0.38 | 0.10 | 0.24 | 0.42 | - | 0.26 | 0.06 |
