## Supplementary Table 5 for "Purine and pyrimidine synthesis differently affect the strength of the inoculum effect for aminoglycoside and β-lactam antibiotics"

**Supplementary Table 5: P values and the exact number of biological replicates (*n*) for Fig. 4.** P values for growth rate and [ATP] were determined using a two-tailed t-test. P values for ATP and growth rate from two-tailed t-test. ΔMIC from a one-tailed t-test relative to nitrogenous base control. We used a one-tailed t-test as we only asked if ΔMIC could decrease (but not increase); thus, we asked our question in one direction only. strp = streptomycin; carb = carbenicillin.

| **Condition** | **[nitrogenous base] mM** | **% CAA** | ***n* (growth rate)** | **P value (growth rate)** | ***n* ([ATP])** | **P value ([ATP])** | **P value (ΔMIC strp)** | **P value (ΔMIC carb)** |
|---|---|---|---|---|---|---|---|---|
| No nitrogenous bases (control) | 0 | 0.1 | 5 | 1.000 | 3 | 1.000 | 0.5 | 0.5 |
| | | 0.5 | 6 | | 3 | | | |
| | | 1 | 6 | | 3 | | | |
| Adenine | 5 | 0.1 | 5 | 0.069 | 4 | 0.032 | 0.385 | 0.051 |
| | | 0.5 | 6 | | 4 | | | |
| | | 1 | 6 | | 4 | | | |
| Cytosine | 10 | 0.1 | 5 | 0.205 | 4 | 0.446 | 0.067 | 0.10 |
| | | 0.5 | 5 | | 4 | | | |
| | | 1 | 5 | | 4 | | | |
| Thymine | 10 | 0.1 | 5 | 0.231 | 4 | 0.669 | 0.041 | 0.037 |
| | | 0.5 | 4 | | 4 | | | |
| | | 1 | 6 | | 4 | | | |
| Uracil | 10 | 0.1 | 5 | 0.520 | 4 | 0.130 | 0.283 | 0.28 |
| | | 0.5 | 6 | | 4 | | | |
| | | 1 | 5 | | 4 | | | |
