## Supplementary Table 6 for "Purine and pyrimidine synthesis differently affect the strength of the inoculum effect for aminoglycoside and β-lactam antibiotics"

**Supplementary Table 6: P values and exact number of biological replicates (*n*) for knockout strains in Fig. 5.** P values were determined using a two-tailed t-test and compared to wildtype strain.

| Strain | [bases] μM | *n* (growth rate) | P value (growth rate) | *n* ([ATP]) | P value ([ATP]) | P value (ΔMIC - strp) | P value (MIC - carb) |
|---|---|---|---|---|---|---|---|
| wildtype (*wt*) | 1 | 7 | | 3 | | | |
| | 4 | 16 | | 3 | | | |
| | 7 | 13 | 1 | 3 | 1 | 1 | 1 |
| *ΔpyrC* | 1 | 13 | | 3 | | | |
| | 4 | 6 | | 3 | | | |
| | 7 | 5 | < 0.0001 | 3 | <0.001 | 0.038 | 0.025 |
| *ΔpurK* | 1 | 19 | | 3 | | | |
| | 4 | 11 | | 3 | | | |
| | 7 | 10 | < 0.0001 | 3 | 0.010 | 0.184 | 0.004 |
