## Supplementary Table 7 for "Purine and pyrimidine synthesis differently affect the strength of the inoculum effect for aminoglycoside and β-lactam antibiotics"

**Supplementary Table 7: Average residual values for growth curve fitting to determine growth rates for knockout strains.** *wt* = wildtype.

| Strain | Base | Biological replicate | | | | | | | | | | | | | | | | | | | | Average Residual | SEM | Overall Average | Overall SEM |
|---|---|---|---|---|---|---|---|---|---|---|---|---|---|---|---|---|---|---|---|---|---|---|---|---|---|
| | 1 | 0.09 | 0.15 | 0.16 | 0.15 | 0.17 | 0.15 | 0.22 | | | | | | | | | | | | | | 0.16 | 0.01 | | |
| | 4 | 0.30 | 0.16 | 0.18 | 0.16 | 0.15 | 0.14 | 0.11 | 0.10 | 0.17 | 0.19 | 0.17 | 0.15 | 0.16 | 0.16 | 0.17 | 0.31 | | | | | 0.17 | 0.01 | | |
| *wt* | 7 | 0.18 | 0.12 | 0.38 | 0.19 | 0.39 | 0.20 | 0.07 | 0.09 | 0.42 | 0.50 | 0.31 | 0.16 | 0.13 | | | | | | | | 0.24 | 0.04 | 0.19 | 0.03 |
| | 1 | 0.03 | 0.19 | 0.25 | 0.06 | 0.05 | 0.06 | 0.06 | 0.16 | 0.21 | 0.27 | 0.24 | 0.11 | 0.18 | | | | | | | | 0.14 | 0.02 | | |
| | 4 | 0.23 | 0.06 | 0.16 | 0.25 | 0.40 | 0.30 | | | | | | | | | | | | | | | 0.24 | 0.04 | | |
| Δ*pyrC* | 7 | 1.31 | 1.97 | 1.94 | 1.59 | 1.55 | | | | | | | | | | | | | | | | 1.67 | 0.11 | 0.49 | 0.10 |
| | 1 | 0.12 | 0.11 | 0.10 | 0.24 | 0.25 | 0.13 | 0.15 | 0.13 | 0.10 | 0.10 | 0.07 | 0.07 | 0.05 | 0.05 | 0.12 | 0.01 | 0.07 | 0.07 | 0.06 | 0.07 | 0.10 | 0.01 | | |
| | 4 | 0.45 | 0.01 | 0.27 | 0.37 | 0.12 | 0.11 | 0.03 | 0.03 | 0.22 | 0.11 | 0.14 | | | | | | | | | | 0.17 | 0.04 | | |
| Δ*purK* | 7 | 0.34 | 0.18 | 0.26 | 0.28 | 0.39 | 0.48 | 0.42 | 0.27 | 0.43 | 0.31 | | | | | | | | | | | 0.34 | 0.03 | 0.18 | 0.03 |
