## Supplementary Table 8 for "Purine and pyrimidine synthesis differently affect the strength of the inoculum effect for aminoglycoside and β-lactam antibiotics"

**Supplementary Table 8: Average residual values for growth curve fitting to determine growth rates in Fig. 6.**

| Growth condition | Nitrogenous base | Biological replicate | | | | | | Average | SEM |
|---|---|---|---|---|---|---|---|---|---|
| 1mM IMP | No bases | 0.02 | 0.01 | 0.17 | 0.02 | 0.08 | | 0.06 | 0.03 |
| | 5 mM adenine | 0.15 | 0.11 | 0.10 | 0.07 | 0.12 | 0.04 | 0.10 | 0.01 |
| | 10 mM cytosine | 0.12 | 0.14 | 0.25 | 0.19 | 0.16 | | 0.17 | 0.02 |
| 0.05 µg/mL 6-MP | No bases | 0.10 | 0.15 | 0.15 | 0.14 | 0.17 | 0.17 | 0.15 | 0.01 |
| | 5 mM adenine | 0.06 | 0.13 | 0.07 | 0.06 | 0.05 | 0.01 | 0.07 | 0.01 |
| | 10 mM cytosine | 0.10 | 0.12 | 0.09 | 0.21 | 0.17 | 0.10 | 0.13 | 0.02 |
