## Supplementary Table 9 for "Purine and pyrimidine synthesis differently affect the strength of the inoculum effect for aminoglycoside and β-lactam antibiotics"

**Supplementary Table 9: P values and the exact number of biological replicates (*n*) for experiments in Fig. 6.** P values were determined using a two-tailed t-test and compared to wildtype strain and without inhibitor and nitrogenous base supplementation.

| Growth condition | Nitrogenous base | Biological replicates (*n*) for growth rate | P values for growth rate | Biological replicates (*n*) for [ATP] | P values for [ATP] |
|---|---|---|---|---|---|
| 1 mM IMP | No bases | 5 | 0.026 | 6 | 0.078 |
| | 5 mM adenine | 6 | < 0.0001 | 6 | < 0.0001 |
| | 10 mM cytosine | 5 | < 0.0001 | 6 | 0.0007 |
| 0.05 µg/mL 6-MP | No bases | 6 | 0.32 | 6 | 0.031 |
| | 5 mM adenine | 6 | < 0.0001 | 6 | < 0.0001 |
| | 10 mM cytosine | 6 | 0.923 | 6 | 0.002 |

**Supplementary Table 10: P values and the exact number of biological replicates (*n*) for MIC experiments with kanamycin.** P values were determined using a two-tailed t-test and compared to wildtype strain and without inhibitor/nitrogenous base supplementation.

| Inhibitor | Nitrogenous base | Biological replicates (*n*) for growth rate | P value (ΔMIC) |
|---|---|---|---|
| none | none | 5 | 1 |
| 6-MP | none | 5 | 0.02 |
| | 5 mM adenine | 5 | <0.001 |
| | 10 mM cytosine | 6 | 0.876 |
| IMP | none | 6 | 0.001 |
| | 5 mM adenine | 6 | 0.003 |
| | 10 mM cytosine | 6 | 0.184 |
